# High-speed 3D Imaging with 25-Camera Multifocus Microscope

**DOI:** 10.1101/2024.09.23.614351

**Authors:** Eduardo Hirata Miyasaki, Antone A. Bajor, Gustav M. Pettersson, Maximilian L. Senftleben, Kaitlyn E. Fouke, Thomas G.W. Graham, Demis D. John, Jennifer R. Morgan, Gal Haspel, Sara Abrahamsson

## Abstract

We here report an aberration-corrected 25-plane camera array Multifocus microscope (M25) for high-speed, high-resolution wide-field optical microscopy in three spatial dimensions (3D). We demonstrate live imaging of 25-plane 3D volumes of up to 180×180×50um at >100 volumes per second. 3D data is recorded simultaneously by an array of 25 small, sensitive, synchronized machine-vision cameras. M25 employs aberration-corrected Multifocus microscopy—an optical method where diffractive Fourier optics are used for multiplexing and refocusing light— with a simplified design for chromatic dispersion correction where a corrective diffractive gratings is placed on each camera in the array. This elegant architecture for chromatic correction will be applicable in a broad range of diffractive imaging applications. M25 is a powerful optical tool for high-speed 3D microscopy in that it allows both non-invasive, label-free bright-field and highly sensitive fluorescence microscopy. We showcase M25 capabilities in 3D particle tracking, bright-field, and fluorescence imaging in *D. melanogaster*, and locomotion and neural activity studies in *C. elegans*.

## 1.0 Introduction

Optical microscopy is an important tool in biological and biomedical research, allowing live, high-resolution visualization of cells, tissues, and small living organisms under physiological conditions. Over the past few decades, fluorescent indicators of neural activity have revolutionized neurobiology by enabling the visualization of neural signaling. Small and transparent model organisms such as the nematode *C. elegans*, the lamprey *P. marinus*, and the fruit fly *D. melanogaster* with genetically expressed fluorescent indicators of neural activity are widely used model organisms in basic research on neural circuit function.^1^

One of the major challenges in live microscopy— particularly in functional neuronal imaging and embryology—is that optical microscopes inherently form two-dimensional (2D) images, while specimens and dynamics of interest are most often three-dimensional (3D). In classical wide-field microscopy, 3D images are formed by the sequential collection of a focus (*z*-) stack of finely sliced 2D lateral (*x,y*) image planes of the specimen, building up the 3D image volume. Refocusing can be performed by varying the distance between the specimen and the objective using a fast piezo *z*-stage or by a remote focusing method.^2^ Confocal microscopes scan a laser laterally and axially^3^ to form a 3D image. However, even with high-speed hardware, the process of sequentially acquiring 3D image data can be prohibitively slow when studying fast and dynamic biological events in the millisecond regime, such as locomotion, transcription, diffusion of biomolecules, or neural circuit activity. This significantly limits our capability to visualize and study crucial biological processes in 3D.^4^

Light sheet fluorescence microscopy (LSFM) has recently provided a faster scanning solution to this challenge. LSFM employs a thin sheet of excitation light to selectively illuminate and excite a single plane of the specimen, thereby reducing phototoxicity and photobleaching.^5–7^ However, LSFM and other traditional methods face limitations in spatiotemporal resolution as sequential scanning of the 3D volume is still required.

A few approaches for capturing truly simultaneous wide-field 3D images, such as light-field microscopy, have been developed. Most of these methods suffer from reduced resolution, require advanced and computationally intensive image reconstruction that can yield ambiguous results, or do not form classical 2D images.^8^ (Methods that do not form classical 3D images can be useful for example in particle tracking.^9^) Extended focus microscopy^10, 11^ is another method to cover a deeper 3D volume simultaneously, but suffers from loss of image peak intensity and resolution.

Aberration-corrected multifocus microscopy (MFM)^12^ provides a unique solution to the simultaneous 3D imaging problem. Here, the image beam from a wide-field microscope is simultaneously multiplexed and refocused by a specially designed diffractive optical element—a multifocus grating (MFG)—to project an entire 3D focus stack from the specimen volume onto a single image plane. The stack of refocused 2D planes is laid out in an array on a digital camera sensor and captured in a single shot without mechanical movement of any kind. Since the MFG is placed in the Fourier plane of the microscope, the entire spatial frequency support (optical transfer function^13^) of the objective is maintained. Data processing is minimal; individual focal planes are cropped out and laterally (*x,y*) registered on top of each other to form the 3D image volume. Successive focal planes are separated by an evenly spaced focus step Δz, which can be tailored to the application at hand in the MFG design.

High-resolution (high Numerical Aperture) microscope objective lenses are designed to provide aberration-free images only from their primary focal plane. Severe aberrations—dominated by depth-induced spherical aberration and increasing in magnitude with refocusing depth—are introduced when the microscope objective lens is used away from its nominal front focal plane.^2^ MFM employs an aberration-free refocusing function, enabled by the use of diffractive Fourier optics, to eliminate this effect. A geometrical distortion—calculated according to the Abbe sine condition^13^—of the nominally periodic MFG grating pattern corrects for the depth-induced aberrations that would otherwise arise^12^ when the microscope objective is used to image a plane that is not at its primary front focal plane. This correction is crucial, as significant spherical aberration would otherwise be introduced after a defocus of a couple of micrometers, deteriorating both image resolution and sensitivity.

In addition to optimizing resolution and image peak intensity, light efficiency is another critical factor in high-speed biomicroscopy. Acquisition speed often becomes effectively limited by signal strength and light dose. Therefore, the diffractive optics components of MFM have been optimized for light efficiency. In a previous publication, we describe a process of nanofabrication of fused silica phase grating MFGs with an excellent quantum efficiency of up to 89%.^14^

While diffractive Fourier optics enable highly efficient beam multiplexing and aberration-free refocusing through a deep imaging volume, the effect of chromatic dispersion—which is inherent to diffractive optical elements—must also be addressed to maintain optimal performance in broadband (color) imaging.^13^ While not necessary in brightfield imaging, this is crucial in fluorescence microscopy on sensitive living specimens. Even a single-color fluorophore is not monochromatic but is typically imaged across a wavelength spectrum of tens of nanometers. Unfortunately, MFM systems for broadband imaging are complex since they contain an optical chromatic dispersion correction module (CCM). The CCM removes the chromatic dispersion introduced by the MFG. The classical MFM employs a CCM consisting of a multifaceted prism and a multi-paneled blazed diffractive grating combination, which provides excellent imaging performance but requires custom optical manufacturing and meticulous alignment.^12^

The original MFM systems were designed to capture up to nine focal planes simultaneously on a single camera sensor.^12^ This is an excellent *z*-sampling range for studying fast dynamical events in smaller specimen volumes, such as transcription inside the cell nucleus^14,15^.^16^ However, high-speed, high-resolution imaging of larger and deeper sample volumes is also desired.

Although using the original MFM chromatic correction design for an arbitrary number of image planes is conceptually simple, manufacturing cost, and complexity increase with the number of planes. Therefore, MFM imaging systems that capture more than nine simultaneous focal planes have been implemented without chromatic correction and have thus not been sensitive enough for live fluorescence microscopy. Even in monochromatic imaging, the use of a single camera to capture 25 image planes has limitations in terms of spatio-temporal sampling resolution.

We have developed an aberration-corrected multifocus 25-plane camera-array (M25) microscope to overcome these limitations. M25 captures simultaneous 3D volumes of 25 aberration-corrected image planes on an array of synchronized digital cameras at over 100 volumes per second (VPS). This powerful instrument employs a radically simplified optical design for chromatic dispersion correction, using pairs of diffractive gratings. M25 is demonstrated in fluorescent and brightfield microscopy of whole, living animals through an up to 50 µm deep volume, to visualize fast, dynamic biological processes in their native 3D contexts. This method for capturing Fig 3D dynamics without sequential scanning represents an advance in the field of live 3D imaging.

## 2.0 Results

Here, we present the design, deployment, optical performance, and live imaging capability of the aberration-corrected multifocus 25-plane camera-array microscope (M25). This apparatus (Figure 1) enables deep, sensitive, simultaneous high-speed 3D imaging of living samples in bright-field transmission and fluorescence microscopy through an extended image volume. M25 employs a new, simple, and powerful method for chromatic dispersion correction with an architecture of paired diffractive phase gratings on a multi-camera array (Figure 2). This architecture could also be used for other imaging modalities, including hyperspectral imaging and spectroscopy applications, that employ image multiplexing. The optical design thus has the potential to open up the use of diffractive optics in broadband imaging in a broad range of applications. We demonstrate M25 imaging in pilot studies of the model organisms *D. melanogaster* (Figure 4) and *C. elegans* (Figure 5,6.7) and *P. marinus* (Figure S8).

**Figure 1.**
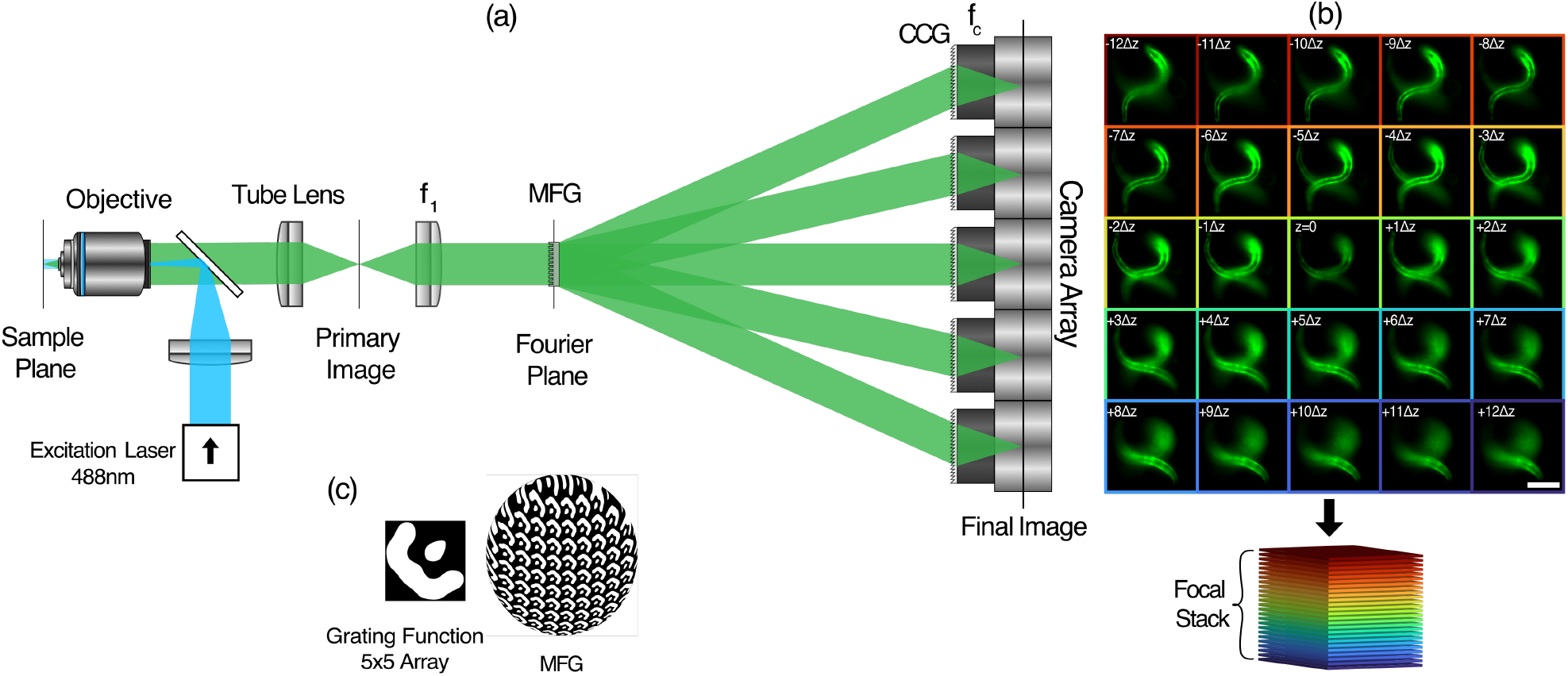
Aberration-corrected 25-plane camera-array multifocus microscope (M25). (a) Beam path of the M25 microscope, connected to the side-port of a wide-field microscope that consists of the fluorescence excitation, an objective, tube lens, and a dichroic mirror. The relay lens *f*_1_ creates a conjugate Fourier plane where the 25-plane multifocus grating (MFG) is placed. The MFG multiplexes and refocuses the image beam to form a simultaneous focus stack. Each 2D focal plane of the stack is recorded on a digital camera equipped with a blazed chromatic correction grating (CCG) and camera lens *f*_*c*_. (b) M25 image of 25 focal planes recorded with the camera array, showing two freely moving and twisting fluorescently labeled *C. elegans* imaged at 50 VPS. A 3D render of the nematodes over time can be found in Video S1. Below: focus stack order. (c) 25-plane MFG grating function and conceptual illustration of the MFG pattern distortion for aberration-free refocusing. Scale bar in (b) 75 µm

**Figure 2.**
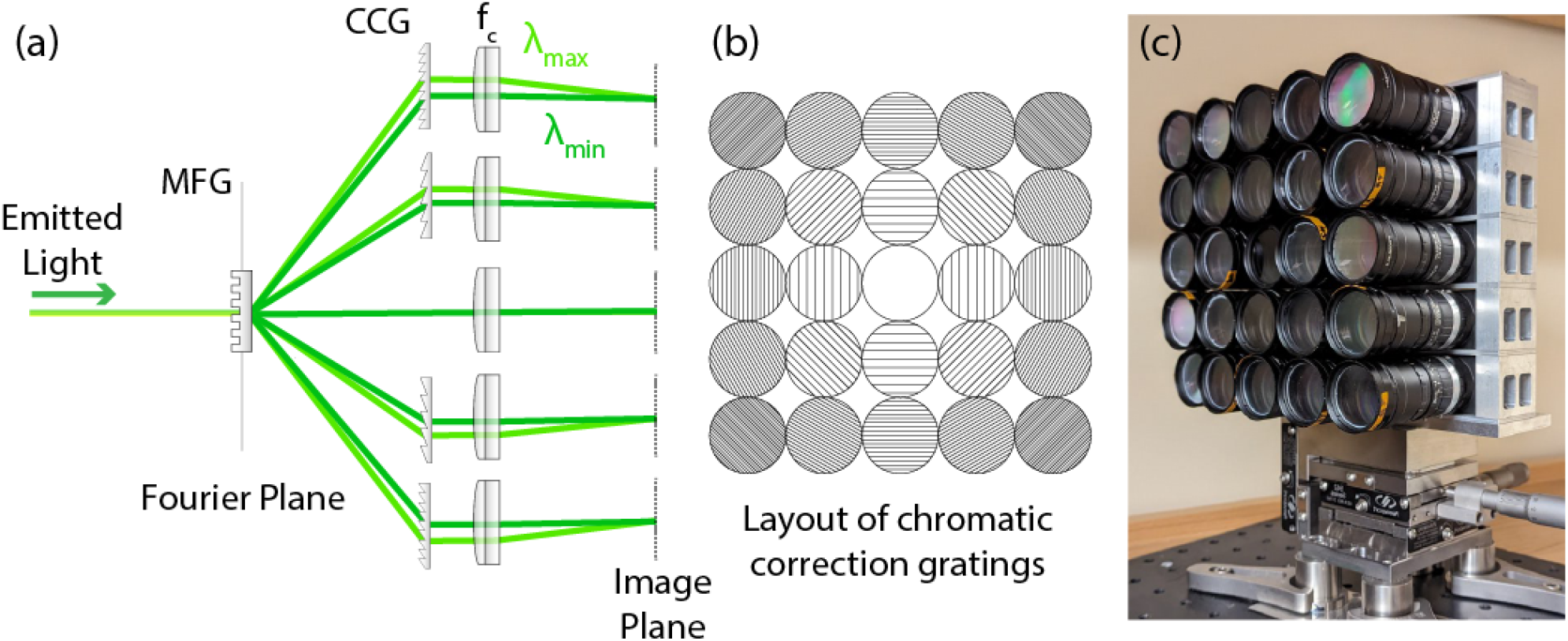
Chromatic Dispersion Correction architecture of M25. (a) Blazed phase gratings, fitted in the filter holders of the camera lenses, remove the chromatic dispersion introduced by the MFG. (b) Chromatic Correction Grating (CCG) arrangement for M25. Different positions have different grating periods to account for the magnitude and orientation of dispersion in the 2D diffraction orders *m*_*xy*_. (c) Photograph of the 5 × 5 camera array, mounted into a custom-built aluminum frame, each camera lens outfitted with a CCG.

**Figure 3.**
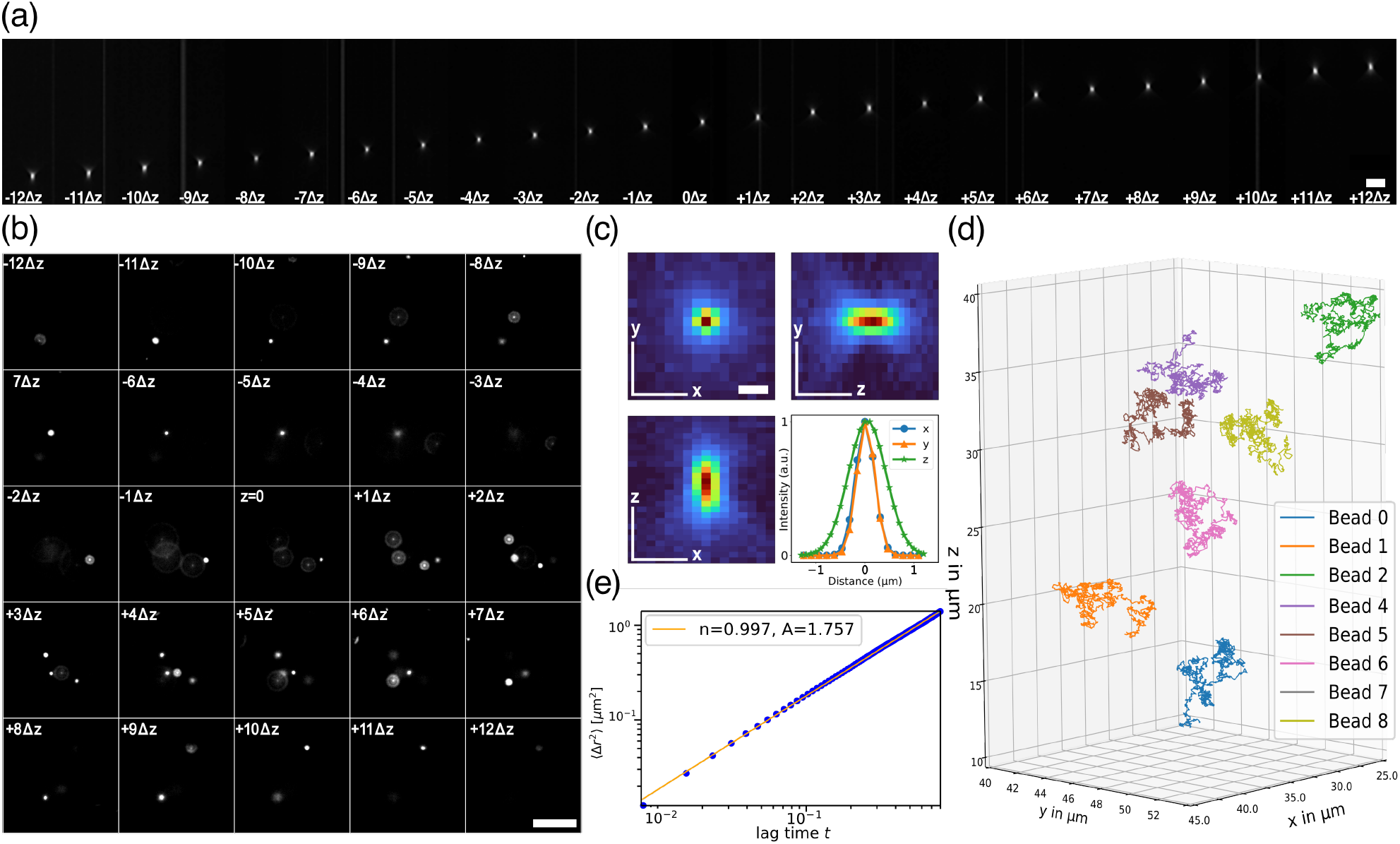
Verification of resolution, aberration-free refocusing, and high-speed image acquisition. a)Point-spread-function (PSF) montage of M25 imaging system using 200 nm fluorescent beads with aberration-free refocusing across a depth of 50 µm. The M25 successive focus step is Δ*z*=2 µm, and the lateral pixel size is 156 nm. The M25 focus stack was recorded with an axial sampling of 100 nm z-steps and has been cropped and re-scaled to display the axial PSF of each focal plane with isotropic voxels. b) Fluorescent beads (1 µm) diffusing in water imaged at 125 VPS to verify M25 hardware control and acquisition software capability. c) Representative PSF (of plane ™8Δ*z* in *a*) showing the FWHM in xy-, xz- and yz-projections together with a Gaussian-fitted FWHM curve for the three axes. The mean FWHM over all planes is x: 365.2 ±57.8 nm, y: 358.8 ±26.4 nm and z: 685.1 ±114.8 nm. d) Video of 3D tracking of beads in Video S2. e) Measured diffusion coefficient of beads in *b*. Scale bar in (a) is 3 µm, (b) is 25 µm and in (c) is 1 µm.

**Figure 4.**
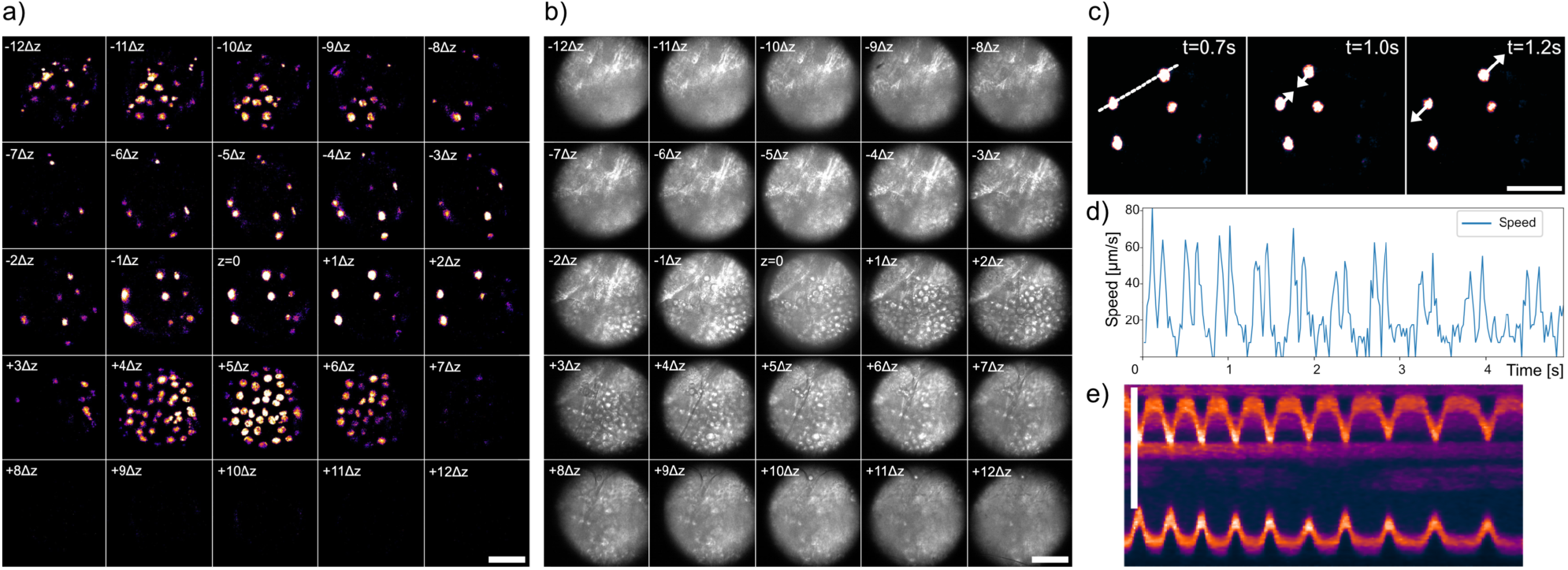
Live fluorescence and brightfield M25 imaging of the heart of transgenic *Drosophila* larva expressing EGFP-tagged histone H2B. a) First time frame of a time series showing cell nuclei labeled with H2B-EGFP over five seconds. b) First time frame of the same cells from *a* in bright-field mode. The fluorescence and brightfield imaging over time can be seen in Video S3. c) Cell contractions at plane +1Δz from *a* at timepoints t=0.7s,1s, and 1.2s. d) Intensity profile along the dotted line in *c*. e) Kymograph showing the speed of tissue contraction over a 5-second time frame. Scale bar 25 µm.

**Figure 5.**
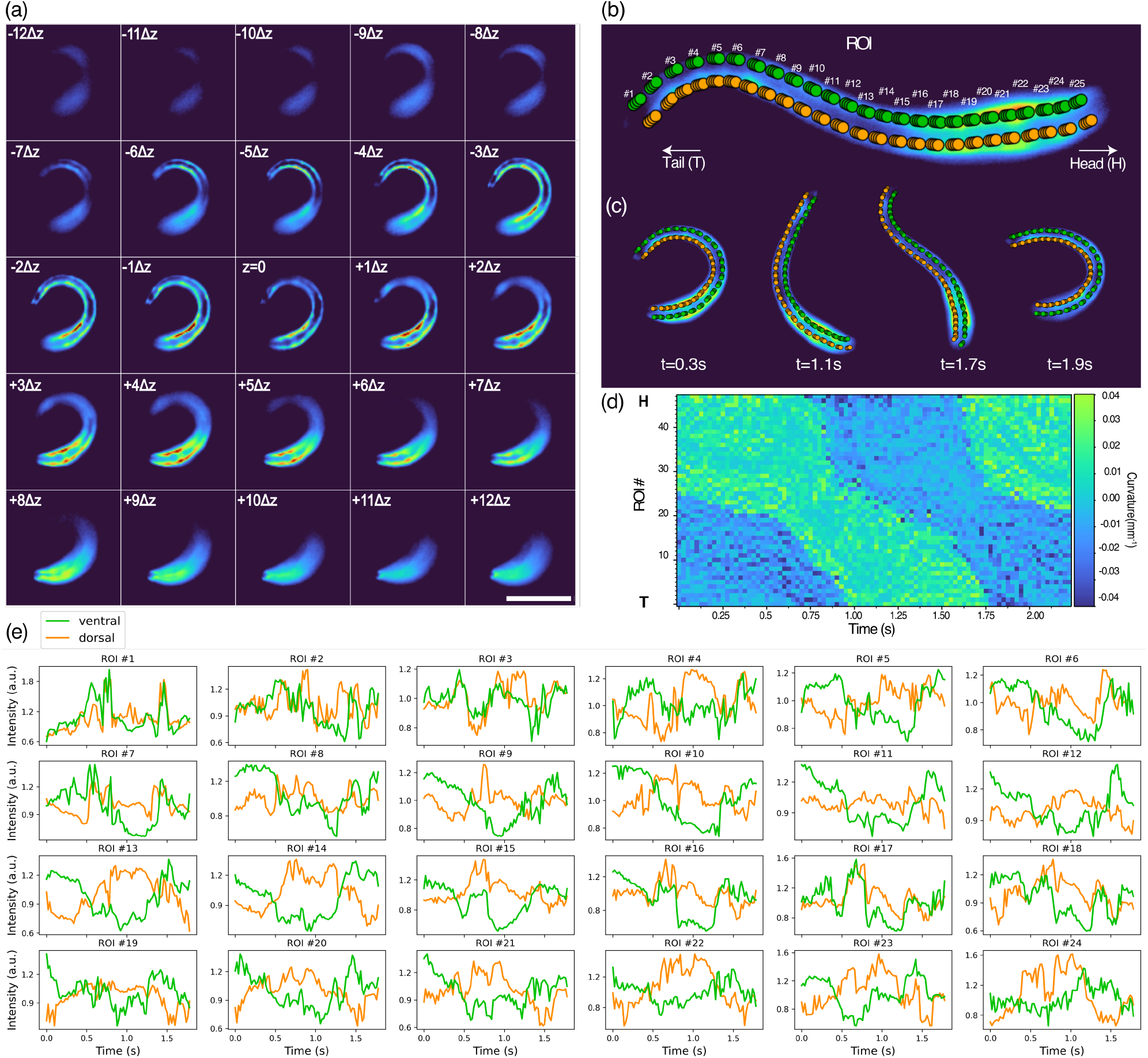
Alternating activation of antagonistic muscles during locomotion in *C. elegans* expressing the fluorescent calcium indicator GCaMP (a) Single time point from Video S4 acquired at 50 VPS (b) ROIs for tracking neural activity during ROI No.0 being the tail and No.49 the head (c) Defined four circular ROIs placed between the skeleton interpolated points to identify muscles individually (d) Poses of the undulating nematode in *a* at time frames *t* = 0.24, 0.86, 1.34, 1.54*s*. (e) Green and blue shaded areas in the kymograph represent the dorsoventral curvature of the animal in *a* where the axis is the length of the animal from tail (T) at the origin and head (H) at the top along time (horizontal axis). e) Fluorescence intensity (calcium indicator GCaMP) of the ROIs in *c* over time showing muscle activity during free movement. Scale bar in (a) is 70 µm.

**Figure 6.**
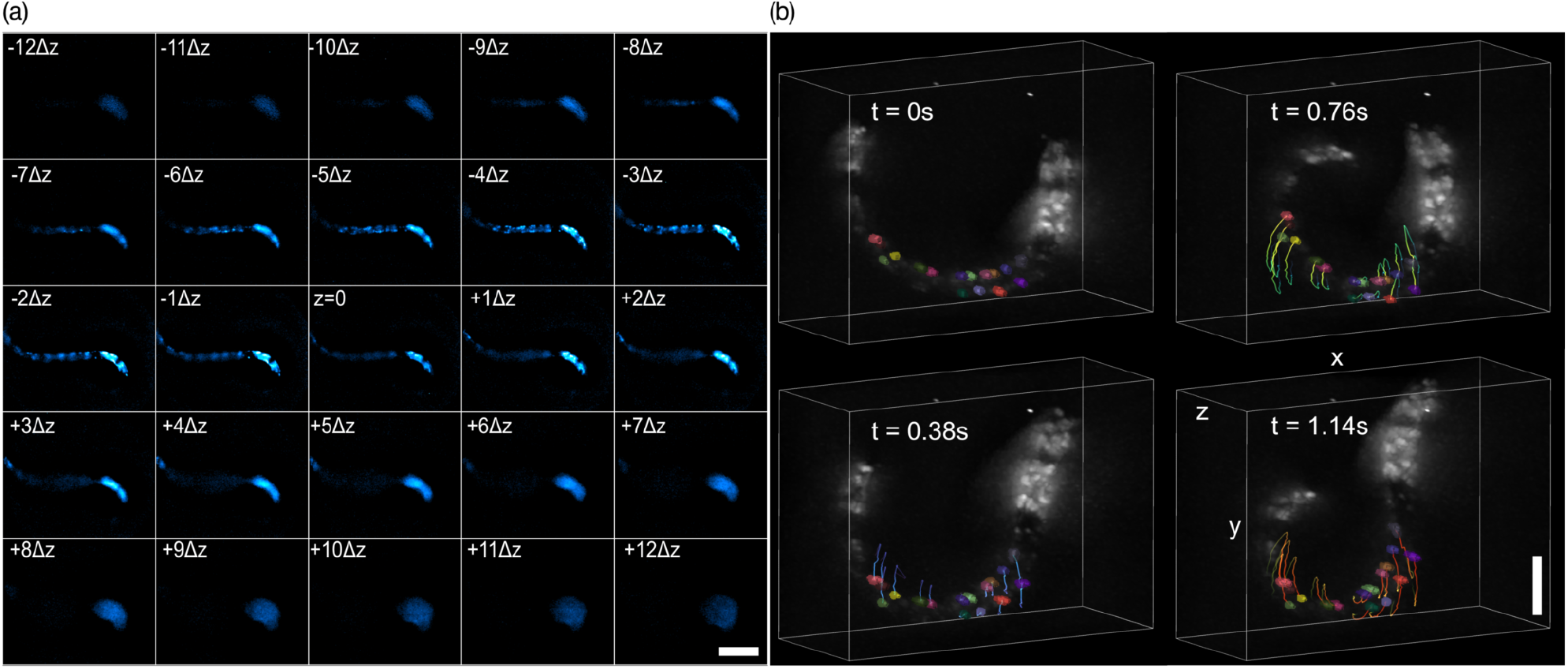
Rapid movement of *C. elegans* and tracking of individual neurons. (a) Raw time frame from Video S5 of a of freely moving animal showing each plane. (b) Tracking of neurons around the midsection of *C. elegans* from (a) over time. The neurons were segmented, then tracked, and finally visualized using Napari. (Video S6) Scale bar in (a) 40 µm and (b) 20 µm.

**Figure 7.**
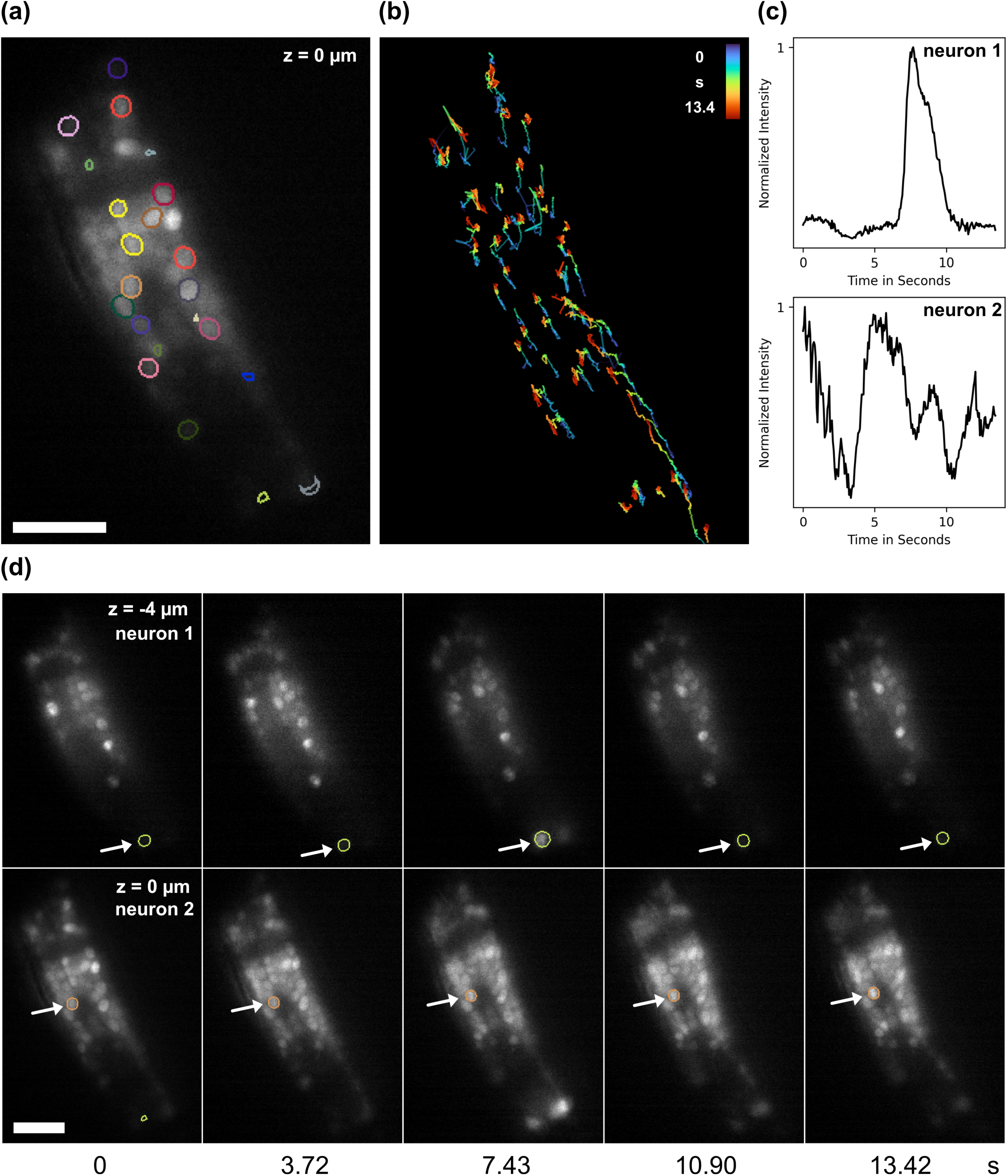
Neural signaling imaging of *C. elegans* head acquired with a smaller M25 field-of-view with a voxel size of 0.156 × 0.156 × 2.0 µm. Neurons were individually labeled in all planes, (a) shows a representative plane. (b) Visualization of spatial neuron dynamics; the individual neurons were linked over time, short linkages were filtered out and then presented with the LUT Turbo for all planes. (c) Normalized intensity profiles of two distinct neurons from two different axial planes, (d), show calcium activity through the fluorescence response of GCaMP over time. The frame rate was 16 VPS. The volumetric imaging over time can be seen in Video S7. Scale bar in (a), (b), and (d) 5 µm.

### 2.1 M25 optical design

The M25 microscope employs multifocus diffractive Fourier optics for aberration-free refocusing.^12^ The image beam from a commercial wide-field microscope is multiplexed by a 25-plane MFG (Figure 1 a) into a 5 × 5 array of images formed by the set of the zeroth, first, and second 2D diffractive orders *m*_*xy*_ (*x, y* : *−* 2 *→* +2) (Figure 1 b). The 25-plane MFG grating function (Figure 1 c) adjusts the energy distribution with optimal light efficiency (≈ 80% quantum efficiency) and evenness between the desired diffractive orders with minimal light loss for highly sensitive imaging.^14^ Each diffracted beam is refocused by the MFG through a geometrical distortion of the grating pattern (Figure 1 c). The distortion function for aberration-free refocusing is calculated according to the Abbe sine condition to correct the out-of-focus phase error of the defocus Δ*z*, as described in a previous publication.^12^ The magnitude of the distortion determines the focus step Δ*z* between successive planes and is adjusted to fit the best compromise between *z-*sampling and volume depth for the application at hand. We have designed and tested 25-plane MFGs with focus steps Δ*z*=1 µm and Δ*z*=2 µm to cover volume depths of 25 µm and 50 µm, respectively. (Data is shown from the systems covering the largest volumes.)

A key feature of the M25 optical design is a powerful and elegant new architecture for chromatic dispersion correction in a camera array(Figure 2.) Diffractive Fourier optics elements provide precise wavefront control at every point in the objective pupil (Fourier) plane but introduce chromatic dispersion. According to the grating equation sin *θ*_*m*_ = *mλ/d*,^13^ the diffraction angle *θ*_*m*_ of each diffractive order *m*_*x,y*_ depends on the grating period *d* and the wavelength *λ*. Strong chromatic dispersion is thus introduced when imaging across a wavelength spectrum wider than a few nanometers. This dispersion problem is a major reason why diffractive optics are not more widely employed in imaging, despite their unique capabilities in beam multiplexing and the exquisite control they offer in shaping the wavefront. Without correction, the effect of chromatic dispersion is a smearing of the image in the radial direction, which reduces both resolution and image peak intensity. In fluorescence microscopy, the fluorescence emission light is typically collected across a wavelength spectrum of approximately 30-50 nm to maximize signal in sensitive living specimens covering the broad emission peaks of fluorescent proteins. Thus, even when imaging a single-color fluorophore such as the green fluorescent protein (GFP), chromatic dispersion must be corrected for optimal performance. While manufacturing a prism-grating combination with 25 facets is conceptually straightforward, it is complex and costly in practice. Additionally, capturing an array of 25 images on a single camera sensor^14^ is sub-optimal, as it limits the lateral sampling resolution. The M25 employs a 5 × 5 camera array to address these two major challenges, with each focal plane captured on a separate camera. In front of each camera lens—except the center camera, which captures the non-diffracted zeroth order beam *m*_0,0_—a blazed grating is mounted with a period and angle adjusted to match the dispersion of each 2D diffractive order *m*_*x,y*_. These blazed gratings, referred to as chromatic correction gratings (CCGs), completely remove the chromatic dispersion introduced by the MFG.

Employing an array of 25 cameras in a single microscope would have been a major investment and undertaking two decades ago, but is today quite practical due to the availability of fast, sensitive, and affordable machine vision cameras. Thanks to their small packaging, we can fit the cameras next to each other in a tight array. Off-the-shelf camera lenses with equally small footprints are then fitted into the camera’s c-mount threads. Each CCG is cut with a laser cutter to the shape of a circular disc to fit directly into the filter holder slots of the camera lens mounts. With this architecture, all 25 beams maintain their separation without needing the prism component of previous MFM systems.

All diffraction optical elements for our prototype M25 instruments were manufactured hands-on by students in our team at the UCSB Nanofabrication facility. It can be noted that it would be straightforward to adjust the layout for a larger or smaller number of focal planes—such as 3 *×* 5 = 15 planes or 7 *×*7 = 49 planes—to optimize the imaging depth for specimen size. The described design architecture can be adapted to enable other diffractive optics imaging systems that use diffraction to multiplex beams.

### 2.2 Performance Evaluation of the M25 Multifocus Microscope

We constructed two M25 microscope systems to evaluate the performance and adaptability of the design. One system was assembled at the Marine Biological Laboratory (MBL) at Woods Hole, and the other at the University of California Santa Cruz (UCSC) Microscopy Center. Both systems were tested for imaging speed, resolution, and their ability to capture 3D images across various sample types. To verify optical performance, we imaged 200 nm Tetraspeck fluorescent beads. These bead images were used to measure the point-spread-functions (PSFs) of the system (Figure 3a). The lateral resolution was determined to be approximately 365 nm on average, with an axial resolution of around 685 nm. These values confirmed that the M25 achieved diffraction-limited resolution across the entire 50 µm imaging depth, demonstrating effective aberration-corrected refocusing in each of the 25 focal planes. The uniformity of resolution across all planes highlights the capability of the M25 to maintain image quality even at different depths, a critical feature for high-resolution 3D imaging. The PSF measurements can be used as fiducials to register and align (*x*- and *y*translation) the 25 focal planes from the individual cameras.

To further test the system’s performance, we imaged 1 µm fluorescent beads diffusing in water. The M25 tracked these beads in 3D at a high frame rate of 125 VPS (Figure 3b-d and Video S2), successfully capturing their movement without any dropped frames or motion artifacts, using our custom acquisition engine^17^ and GUI.^18^ We calculated the diffusion coefficient by analyzing the bead trajectories, which matched theoretical expectations. This demonstrated the system’s ability to perform high-speed 3D particle tracking, validating its precision in capturing dynamic events.

### 2.3 Fluorescence and bright-field imaging of *D. melanogaster*

On the M25 microscope at UCSC, we explored multimodal imaging of *Drosophila* larvae. This instrument covered an imaging volume of 50 *×*50 *×*50 µm and performed both fluorescence and bright-field transmission (label-free) microscopy. For bright-field imaging, the microscope was modified by adding a narrow wavelength bandwidth filter after the white lamp in the trans-illumination light path.

We conducted imaging experiments on *D. melanogaster* larvae expressing EGFP-tagged histone H2B, which labels cell nuclei.^19^ The fluorescence imaging mode (Figure 4a) allowed us to track the nuclei, while the bright-field mode (Figure 4 b) enabled label-free imaging, highlighting the structural details and movements of the larvae. Simultaneous 3D imaging of the beating heart of a larva embedded in low melt agarose is shown in Video S3. We quantified heart contractions by measuring the displacement between pairs of nuclei (Figure 4c-e). The observed heart rate was approximately two beats per second, which is consistent with previous observations.^20^ Bright-field transmission and fluorescence microscopy each provide their respective advantages. Bright-field microscopy offers detailed structural information and is minimally invasive, allowing for the observation of sensitive living samples, such as developing embryos, without staining. Fluorescence microscopy, on the other hand, provides high specificity and sensitivity for visualizing proteins or structures within cells. The capability of MFM to perform both bright-field and fluorescence microscopy enables comprehensive imaging, where structural context from bright-field can be integrated with specific molecular insights from fluorescence. This capacity uniquely poises multifocus microscopy within high-speed 3D imaging microscopy methods, most of which can only be used in fluorescence mode, for a wide range of biological applications.

### 2.4 Functional neuronal imaging in *C. elegans*

We explored functional neuronal imaging in *C. elegans* using the M25 microscope at MBL Woods Hole(Figure 5). This instrument covered a volume of 180 ×180×50 µm with 50 VPS cameras.

We measured coordinated muscle activity in *C. elegans* expressing GCaMP2 to study locomotion and track calcium dynamics in muscle subgroups derived from the 95 body wall muscle quadrants along the anterior-posterior axis.^21^ To ensure natural locomotion, animals were placed in an NGM buffer solution, avoiding immobilization or the use of viscous media, which could interfere with typical movement (see Methods, 5.4). This experimental setup allowed for continuous and unimpeded locomotion, enabling the precise measurement of calcium activity in specific muscle subgroups during movement. Our approach provided detailed insights into the dynamics of locomotion, allowing us to effectively analyze muscle activity patterns in freely moving C. elegans.

To propel itself, the nematode uses dorsoventral bending of its body, which includes forward motion, backward locomotion, dwelling, and quiescence.^22, 23^ This motion, controlled by alternating contractions of antagonistic muscles along the body,^24^ requires high frame rate 3D imaging of freely moving animals to allow us to capture activity patterns generated by different motor neurons at varying frequencies.^25^ Observing young, freely moving animals within the field of view (FOV) provided valuable insights into their natural locomotion behaviors, demonstrating the effectiveness of the M25 system in capturing dynamic biological processes in real-time. These patterns can be used to refine existing computational models^26^ and to study complex movements like roll maneuvers for active reorientation.^27^

Next, we imaged *C. elegans* OH1625 strain, that yields a bright panneuronal nuclear GCaMP6s expression and allows tracking of individual neurons over time (Figure 6). We acquired a time-series with our M25 system at 43 volumes per second with a FOV of 180 × 180 × 50 µm^3^, big enough to fit the whole moving animal. From this timelapse, the neurons in the central part of the animal were segemented using *StarDist3D*,^28^ and tracked with *ultrack*^29^ to link neurons over time. Similarly to Figure 5, the animal could move freely within the observed volume (Figure 6b and in Videos S5 and S6), while individual neurons were tracked over 20 µm.

We then imaged the head of an OH1625 *C. elegans* animal with our higher magnification M25 setup at 16 volumes per second (VPS) to quantitatively observe individual neurons over time (Figure 7). Similarly, we used StarDist3D^28^ to segment the neurons in all planes. Then, TrackPy^30^ was employed to spatially link the neurons over time and to analyze the calcium activity behavior. The tracks were filtered (existing in more than 50% of the time frame captured) and show the general movement of the animal over time (Figure 7). We chose two representative neurons from two different z-planes, axially 4 µm apart, and plotted their normalized intensity over time (Figure 7c-d). Neuron 1 exhibits a strong calcium peak at around 7 seconds, while neuron 2 shows multiple smaller peaks over the recorded time frame.

## 3.0 Discussion

High-speed, high-resolution 3D imaging is a central challenge in the field of biological microscopy and a classical problem in the field of optics.^4^ Live 3D microscopy of sensitive, living samples requires careful compromises to capture and quantify critical information at sufficient signal and acquisition speed and without photo-damaging the specimen, especially in fluorescence microscopy. Major trade-offs in live 3D microscopy are acquisition speed, size of imaging volume, light dose, and sensitivity (photon collection efficiency).

Different types of optical imaging systems have advantages and disadvantages. Confocal and light-sheet microscopes, as opposed to wide-field microscopes, have the advantage of optical sectioning and can provide high signal contrast also in densely labeled specimens.^31^ Lightsheet microscopes have the additional advantage of reduced photobleaching, selectively illuminating and imaging one specimen plane at the time. With piezoelectric stages and galvanic mirrors, focus stacks can be acquired at high speed with fast refocusing systems,^32^ remote refocusing,^2^ and swept light-sheet.^7, 33, 34^ Still, these strategies rely on sequentially scanning the sample to collect the focal planes that constitute the image volume; the conflict between speed, resolution/sampling, and imaging volume remains, and there is a temporal ambiguity within the imaging volume. Truly simultaneous 3D image acquisition is a challenging and intriguing concept. The first simultaneous wide-field 3D imaging systems using a multiplane concept were constructed with refractive beam splitters. Off-the-shelf or custom-designed beamsplitters can be used to multiplex the microscope image beam onto multiple cameras, which are offset from the primary image plane to create a defocus.^4, 35–37^ These methods have a great advantage compared to the first diffractive optics method used for multiplane imaging^38^ of avoiding the problem of chromatic dispersion. Unfortunately, depth-induced spherical aberration increases in magnitude with the refocusing depth.^2^ In very shallow volumes, imaging quality can still be acceptable. For example, biplane imaging can be used for single-molecule localization microscopy in *z* using two simultaneous focal planes offset by a few hundred nanometers.^39^

MFM systems for monochromatic imaging are relatively simple to build and can be constructed of just a single custom diffractive optic, the MFG, and two relay lenses. The first lens transforms the microscope image to Fourier space, where the MFG is placed, and the second brings the light back to image space.^14^ Chromatic dispersion is negligible when imaging a narrow wavelength bandwidth of a couple of nanometers. For application in monochromatic bright-field transmission microscopy, the trans-illumination light lamp of a commercial microscope can be cut down by a laser line filter. A simple setup like this would be excellent e.g. in embryology research. Monochromatic MFM systems can also be innovatively implemented using a spatial light modulator^40^ instead of a fixed MFG made of glass, to add flexibility,*e*.*g*., allowing tuning of the focus step during operation. However, chromatic dispersion must be corrected to ensure that full resolution and sensitivity are maintained across the color spectrum in MFM. While MFM^12^ imaging systems with chromatic dispersion correction are complex to construct, they are unique in forming truly simultaneous wide-field 3D images without compromising resolution.

The M25 imaging system here demonstrated thus has potential for duplication and commercialization, as it has a significantly simplified chromatic dispersion correction architecture compared to the original MFM. The design is flexible, and the focus step between planes, and the number of focal planes imaged, can be scaled up or down to suit the application by adjusting the MFG grating function and changing the number of cameras used.

An M25 instrument can be constructed from off-the-shelf components with a set of diffraction gratings (MFG and CCGs) that can be manufactured hands-on at a nanofabrication facility (S10). It should be noted that diffractive optics can be made with varying transmission efficiency depending on the choice of materials and manufacturing process (Figure S9). Our method of manufacturing optimally efficient diffractive phase gratings of fused silica can be duplicated by students or staff in an open nanofabrication user facility. Maintaining light efficiency is crucial for sensitive live imaging to avoid photodamage and to attain high acquisition speed. We have previously provided extensive resources for designing and manufacturing of MFM diffractive optics components.^14^ We also provide resources for open-source hardware control and a user interface for running a large number of fast cameras simultaneously^17^.^18^

To build a multifocus microscope or a chromatic dispersion correction camera array for custom applications, additional resources such as the MATLAB code for designing MFGs, and recipes for nanofabrication of diffractive optics can be found on GitHub^41^ and general tips on optical alignment in our previous publication.^14^

## 4.0 Conclusions

Optical systems that allow truly simultaneous, “single shot” 3D wide-field imaging—without the need for multiple exposures within imaging volume—remove temporal ambiguity within the imaging volume and simultaneously provide high resolution, high 3D acquisition speed, and large imaging volumes.

MFM optical technology provides simultaneous 3D videos at the full resolution of the optical microscope, to be recorded with a temporal resolution ultimately limited by signal strength or the camera readout speed for a single frame. MFM is also compatible with super-resolution and localization microscopy with excellent performance and can be used in both bright-field, fluorescence, and polarization microscopy.^14, 42–44^ MFM can operate in both fluorescence and transmission light modalities, which is important as label-free methods such as brightfield and polarization imaging can be a minimally invasive imaging modality, for example in embryology^45^ and other highly sensitive applications.

Diffractive optics are powerful and allow unique imaging capability through their ability to multiplex beams and precisely control their wavefronts when chromatic dispersion can be corrected. The chromatic dispersion correction through grating pairs on a camera array demonstrated here has potential for application in a wide variety of multiplexing multi-camera imaging applications beyond that of MFM.

## 5.0 Methods

### 5.1 Datasets

Table 1 compiles the datasets from the M25 at MBL and UCSC used in this paper.

**Table 1.** List of all datasets with the corresponding system that was used for acquisition together with the acquired frame rate.

| Dataset | System | Volumes per second (VPS) | Figure |
| --- | --- | --- | --- |
| *C. elegans* | MBL | 51 | Figure 1 |
| M25 PSF | UCSC | / | Figure 3(a) |
| Dancing beads | UCSC | 125 | Figure 3 (b) |
| *Drosophila* larva | MBL | 56 | Figure 4 |
| *C. elegans* locomotion | MBL | 50 | Figure 5 |
| *C. elegans* neurons 1 | MBL | 43 | Figure 6 |
| *C. elegans* neurons 2 | UCSC | 16 | Figure 7 |

**Table 2.**
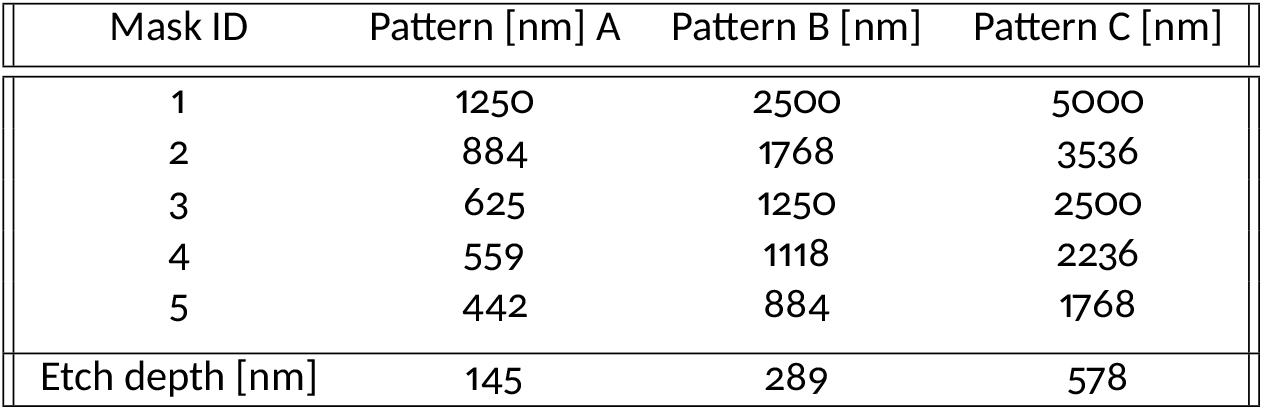
CCG Patterns for all 5 different gratings for 10*µm* period MFG gratings with design wavelength of *λ* = 532 nm. Patterns A, B, and C are shot alphanumerically and etched with their respective to achieve *π* phase shift.

### 5.2 Construction of M25

We developed two M25 microscope setups with varying fields of view (FOV) for different experimental needs. Both systems had no moving parts other than the microscope sample stage and were optimized for 515-540 nm green fluorescence emission, using a Semrock 520/40 bandpass filter. Table 3 summarizes the setups and their corresponding volumetric imaging rates.

**Table 3.**
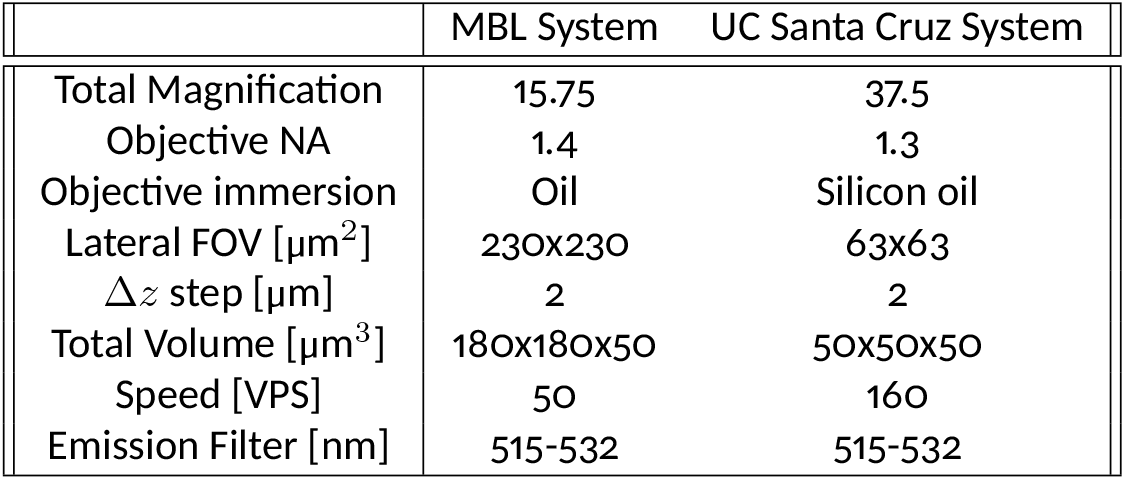
M25 systems imaging specifications for the build at the Marine Biological Laboratory and the UC Santa Cruz imaging facilities.

|  | MBL System | UC Santa Cruz System |
| --- | --- | --- |
| Total Magnification | 15.75 | 37.5 |
| Objective NA | 1.4 | 1.3 |
| Objective immersion | Oil | Silicon oil |
| Lateral FOV [$\mu\text{m}^2$] | 230x230 | 63x63 |
| $\Delta z$ step [$\mu\text{m}$] | 2 | 2 |
| Total Volume [$\mu\text{m}^3$] | 180x180x50 | 50x50x50 |
| Speed [VPS] | 50 | 160 |
| Emission Filter [nm] | 515-532 | 515-532 |

The system at the Marine Biological Laboratory (MBL) was constructed on a Zeiss Axio Observer 7 microscope, equipped with a 63× NA 1.4 oil immersion objective (O1, Zeiss 63x). To create the conjugate Fourier plane where the multifocus grating (MFG) was placed, a relay lens (*f*_1_ = 300 mm, Newport PAC082) was used (Figure S11a,c). The MFG multiplexed the image beam into a 5 × 5 array of focal planes, separated by a 2 µm Δ*z* step. The beams were captured on a camera array consisting of FLIR BFLY-U3-05S2M-CS cameras (6.0 µm pixel size, 50 VPS), with each camera fitted with a chromatic correction grating (CCG) to reverse the chromatic dispersion introduced by the MFG, except for the center camera, which captured the undiffracted zeroth-order beam (Figure S11b). The camera lenses (Fc = 75 mm, Fujinon HF75HA-1B) formed aberration-free images on the camera sensors. The total system magnification was 15.75x. Both the MFG and CCGs were nanofabricated using the recipes detailed in Supplementary Table 2.

The system at UCSC utilized an Olympus IX71 microscope body with a motorized stage (ASI MS-2000) equipped with a 250 µm z-piezo. This system used a 100× silicon oil immersion objective (NA 1.3) and a relay lens (*f*_1_ = 200 mm, Newport PAC087AR.14) to form the conjugate Fourier plane. The MFG, similarly fabricated as in the MBL system, multiplexed the image beam into a 5 × 5 array with either a 1 µm or 2 µm Δ*z* step. Basler Ace 1920-155 cameras (5.86 µm pixel size, 164 VPS) captured the images, with chromatic correction implemented through the same CCG configuration as in the MBL system. Camera lenses (Fc = 75 mm, Tamron M112FM75) were used to form the final aberration-free images. The total magnification of the UCSC system was 37.5x.

### 5.3 Acquisition control

The acquisition control of the M25 system is designed to handle data from 25 cameras simultaneously at high frame rates without dropping frames. This is achieved through a custom acquisition pipeline consisting of three major components: the acquisition engine, the timing control board, and a graphical user interface (GUI).

The M25 system utilizes a custom acquisition pipeline to capture high-resolution 3D images at high frame rates.^17^ The pipeline comprises an acquisition engine, a timing control board, and a graphical user interface (GUI). The acquisition engine simultaneously captures images from 25 cameras at 165 VPS, achieving a data transfer rate of 5.76 GB/s. Data is offloaded using USB 3.0. Each USB has a dedicated host controller via USB PCIe (Neousys cards) to maximize the transfer bandwidth to a RAID0 array of NVME SSDs. Data handling involves writing raw images to a RAM swap buffer and then to disk, ensuring no delays in data transfer. The data is converted to .*raw*, .*tif*, or .*zarr* formats for image processing.

The timing control board or main controller, is a PSoC 5 microcontroller programmed to manage digital blocks and logic for precise timing control of hardware synchronization. It connects to cameras, lasers, stages, and other opto-electronic devices. The main function of the PSoC is to send trigger signals based on GUI-provided parameters (e.g., exposure time, frame rate, capture time). These parameters can be passed through the CLI to the PSoC via USB using *libusb*. Additionally, we developed a Napari plugin that replaces the CLI calls by putting a GUI that wraps and sends the commands via TCP/IP.^18^

### 5.4 Sample preparation

For the verification of spatial resolution, glass coverslips (Carl Zeiss Cover Glasses, High Performance, thickness 0.17 ± 5 µm) were treated with poly-L-lysine (Sigma Aldrich, P8920-100 ML) and coated with 200 nm fluorescent beads (FluoSpheres Carboxylate-Modified Microspheres, F8811, Invitrogen). After that, the coverslips were mounted with ProLong Gold Antifade Mountant (Invitrogen, P36930) and sealed with nail polish. For measuring the diffusion coefficient, 1 µm fluorescent beads (FluoSpheres Carboxylate-Modified Microspheres, F8823, Invitrogen) were suspended in deionized water and added onto a glass coverslip and sealed with nail polish.

#### 5.4.1 C. elegans and D. melanogaster sample preparation

To image the *C. elegans* and *D. melanogaster* samples, were prepared with two different methods to partially immobilize or allow the organism to freely move in space.

For the locomotion and neural activity experiments, the following strains were utilized:

- OH15265 pan neuronal nuclear GCaMp6s: [rab-3::NLS::GCaMP6s + arrd-4:NLS:::GCaMP6s]. Bright pan neuronal nuclear GCaMP6s expression.^46^
- ZW495 body wall muscle GCaMP2: [myo3p::GCamp2 + lin-15(+)]. Transgenic animals are GFP+ in body wall muscle.^47^

The ZW495 samples from Figure 4 and the OH15265 for Figures 6 and 7 were mounted on 2% agarose pads with NGM buffer. A 2 µL suspension containing polystyrene beads (Polystyrene beads 0.1 µm, LB1, Millipore Sigma) with 0.1 µm in diameter at a concentration of approximately 2.5% by volume was added to the pads to partially slow movements following standard methods.^48^

Since the M25 can acquire multiple focal planes simultaneously at high speeds, Figure 3 used 2 µL of 30 µm polystyrene beads (Polystyrene beads 30 µm, 84135, Millipore Sigma)mixed with NGM buffer solution to fully capture moving *C. elegans* in space.^49^

Similarly, the *D. melanogaster* larva sample was mounted on a 2% agarose pad with 2 µl of 30 µm polystyrene beads diluted in water used as spacers to avoid compressing the organism between the coverslip and the pad.

All these samples were covered with (Carl Zeiss Cover Glasses, High Performance, thickness 0.17 ± 5 µm) cover slip sealed with VALAP.

### 5.5 Computational post-processing

The M25 datasets were registered by imaging the same set of beads across the whole volume. We then used the maximum intensity projection and the template matching algorithm to register all the planes with respect to the center camera.

To characterize the performance of this system, the PSFs from the 200 nm beads were analyzed by measuring the full-width half maximum (FHWM) along each projection (XY, YZ, XZ) with the Napari plugin *napari-psf-analysis* by Buchholz.

We measured and tracked the centroids of the 1 µm beads to evaluate the acquisition speed of the M25. The beads were tracked using *TrackPy*, and from the tracks, we calculated the diffusion speed and diffusion coefficient.

The drosophila’s heart beating rate was measured by tracking two cell nuclei using *TrackPy* to get the centroid coordinates and derive the beating speed. We generated the kymograph by tracing a plane across the beating cells using Fiji.^50^

For the *C. elegans* locomotion study, we began by tracking the nematode’s position using active contours and grayscale morphology. We employed the *skan* Python package^51^ to skeletonize the animal’s body and generate equidistant points along both sides of its muscle body. These points were projected orthogonally to the skeleton to define circular regions of interest (ROIs) for measuring calcium activity in the opposing muscle groups. For each frame, we measured the summed intensity of the 4 consecutive ROIs, calculating the fluorescence intensity from the top 80% of the brightest pixels to minimize artifacts from edge hits or muscle deformation. We then computed the median of the means, *F*_80%_(*t*), and normalized the time series as 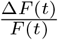. Finally, the 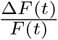 traces were plotted over time, capturing the fast undulatory frequency of the animal to demonstrate forward locomotion in nematodes.^52^

The individual neurons of the *C. elegans* in Figure 6 in the center part of the body were segmented using StarDist-3D.^28^ First, five volumes of the deconvolved time series were manually annotated in 3D using napari, normalized and divided into training and validation data. Next, a stardist model was trained using recommended settings for 400 epochs. The stardist model was then used to predict labels for each neuron in 3D per time frame. Next, Ultrack^29^ was used to link the labels over time. The raw data, the labels, and the tracks were finally visualized using Napari. For the higher magnification dataset of Figure 7 of the head of a *C. elegans*, we used StarDist-3D to label each neuron per focal plane. Then, TrackPy^30^ was used to link the neurons and the normalized intensity was plotted and visualized in napari.

## Declarations

### Author Contributions

SA conceived the M25 instrument. SA and EH designed the M25 instrument. EH, GP, and DJ manufactured the diffractive elements. EH and AB constructed the M25 instrument and built and conceived the data acquisition and hardware control electronics and code. JM, GH, EH, KF, TG and SA conceived and performed the biological experiments. EH and MS performed the data analysis.

### Code availability

The acquisition engine developed for the high-throughput simultaneous acquisition of data with 25 cameras can be found here on our GitHub,^17^ the plugin for acquisition GUI can be found here.^18^

## Acknowledgements

We thank Grant Hartzog, Bill Sullivan, Needhi Bhalla, David States (UCSC) and Ben Abrams (UCSC Life Sciences Microscopy Center) and the MBL Central Microscopy Facility for their support and encouragement and for hosting our instruments. We thank Louie Kerr (MBL), Jim McIlvain (Zeiss), and the entire 2019 Neurobiology Course at the Marine Biological Laboratory in Woods Hole, MA, for their participation in this project. We thank the incomparable Nico Stuurman of Micro-Manager (UCSC) for helpful discussions in high-speed multi-camera synchronization. We thank Brian Thibeault and all the amazing staff at the UCSB Nanofabrication Facility for supporting and training our students in the nanofabrication of optical devices. We thank Mike Stadler (UC Berkeley) who generated the H2B-EGFP *Drosophila* line. Finally, we thank Al Cisneros and the *Sleep* group, who, through works like *Dragonaut*, have provided great inspiration.

## Supplementary Materials

**video S1.**
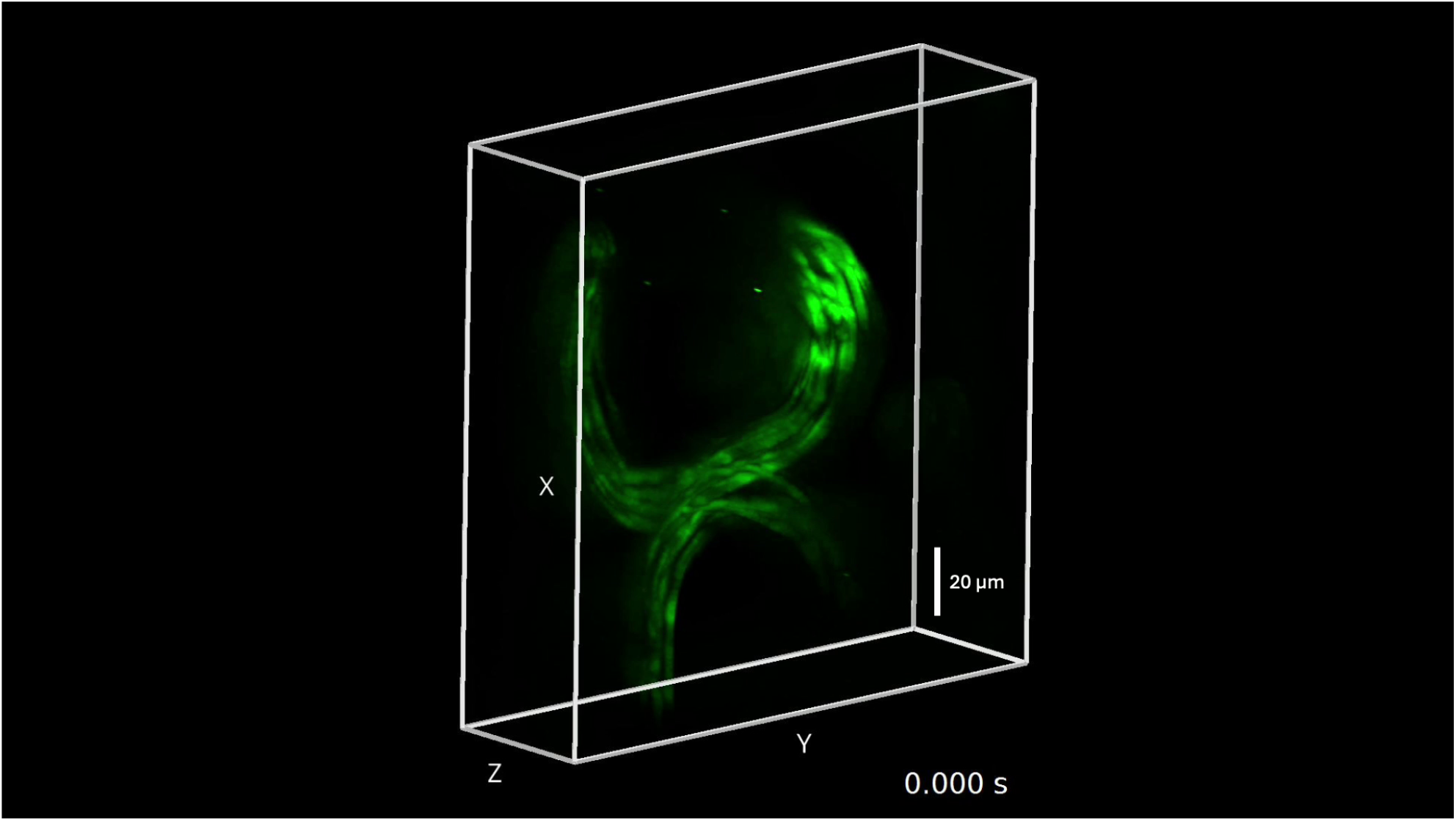
*C. elegans* moving inside the FOV.

**video S2.**
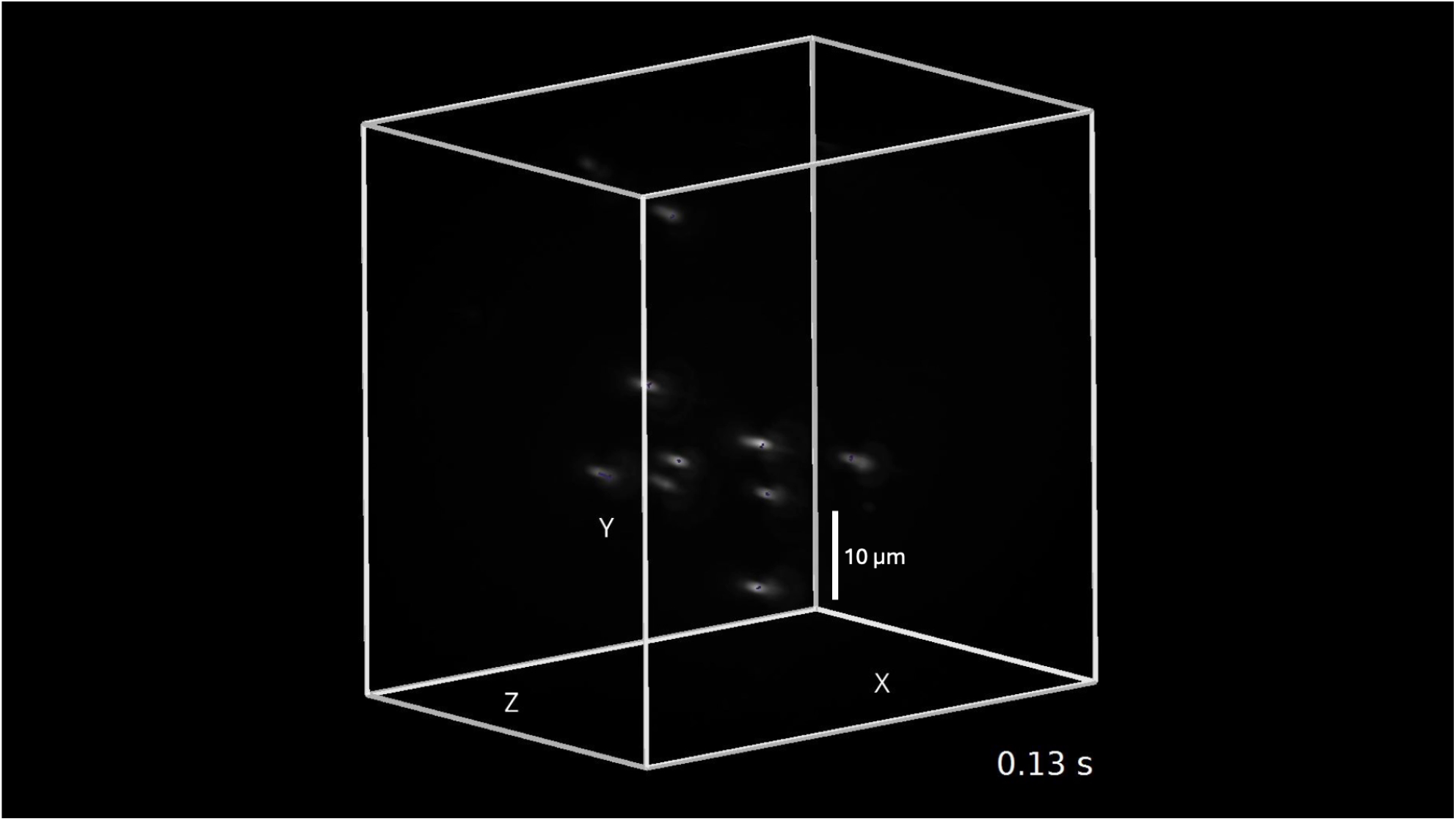
Tracking of 1 µm beads in water.

**video S3.**
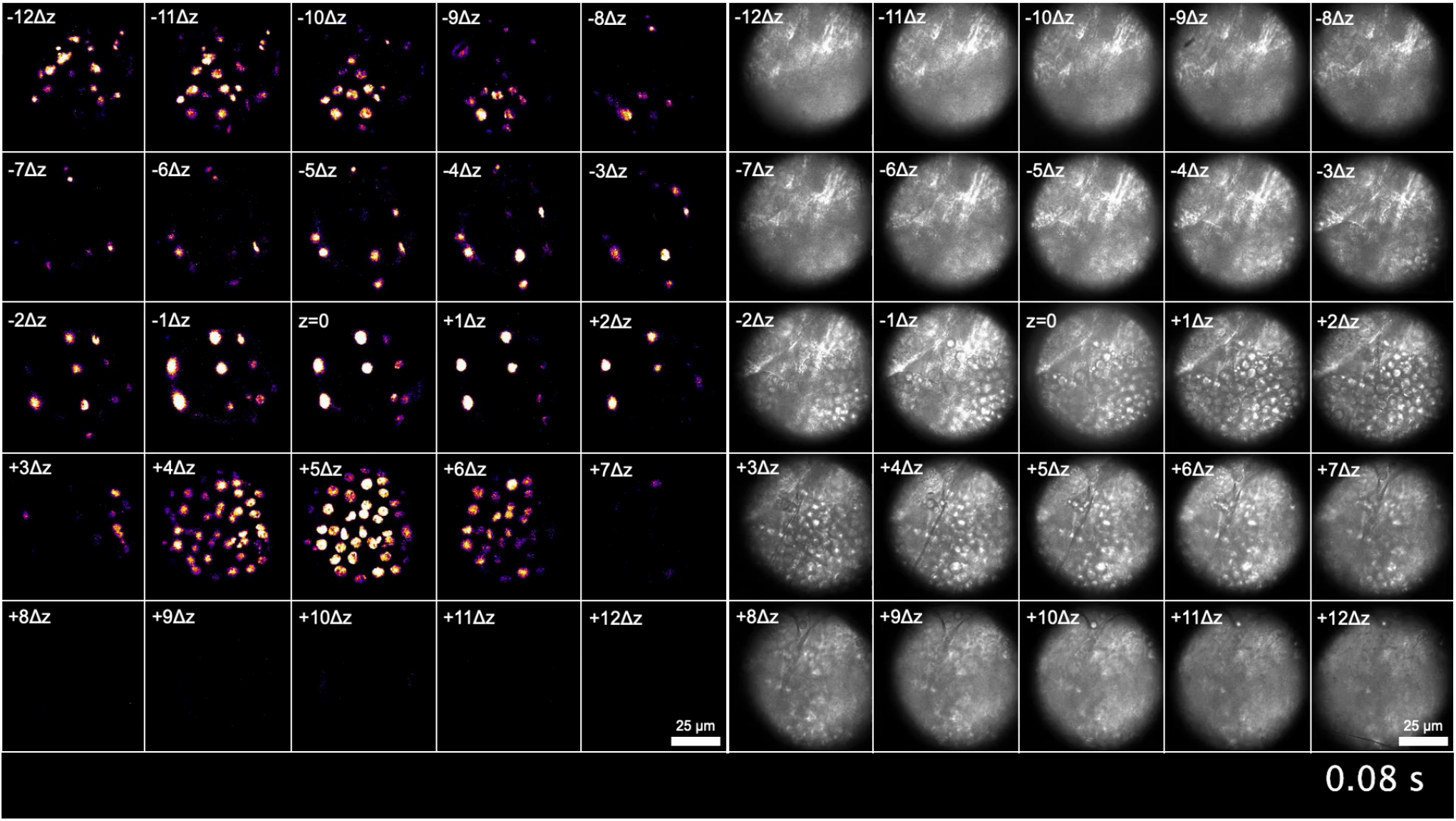
M25 5×5 multiplane montage imaging Drosophila larva H2B using fluorescence and bright field modalities.

**video S4.**
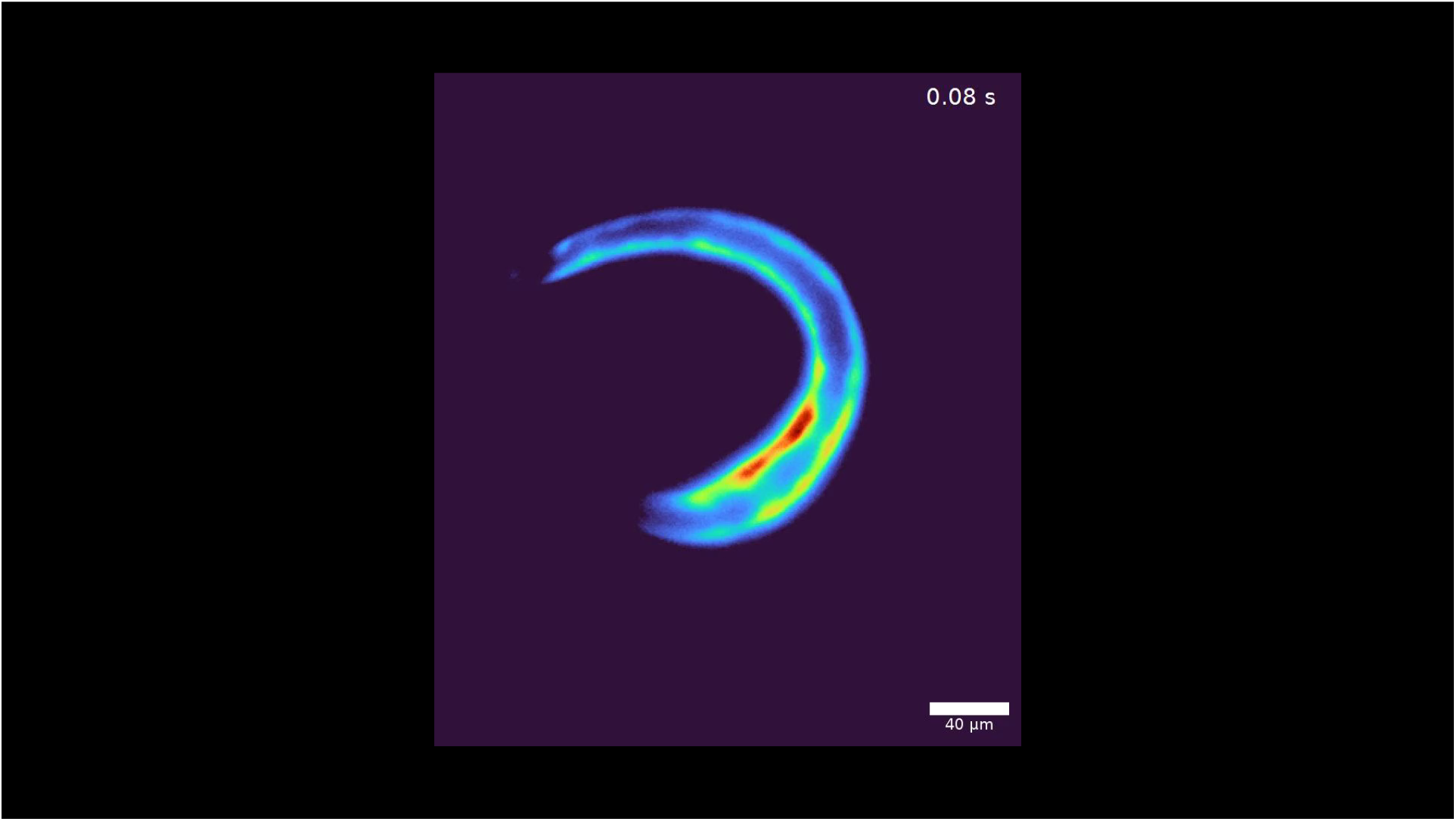
*C. elegans* (ZW495) freely moving.

**video S5.**
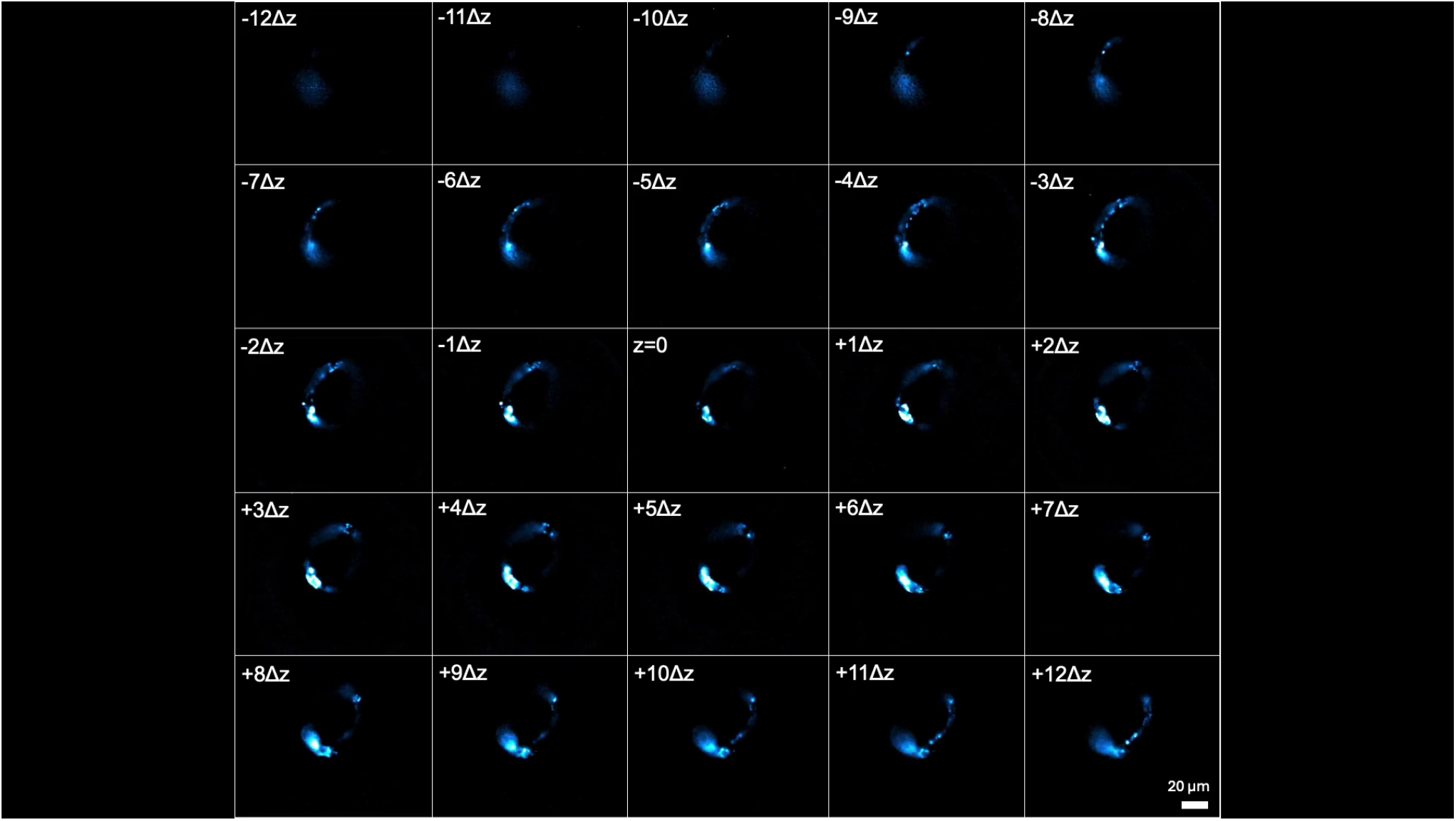
M25 multi-plane montage showing the *C. elegans* (OH1625 strain) freely moving.

**video S6.**
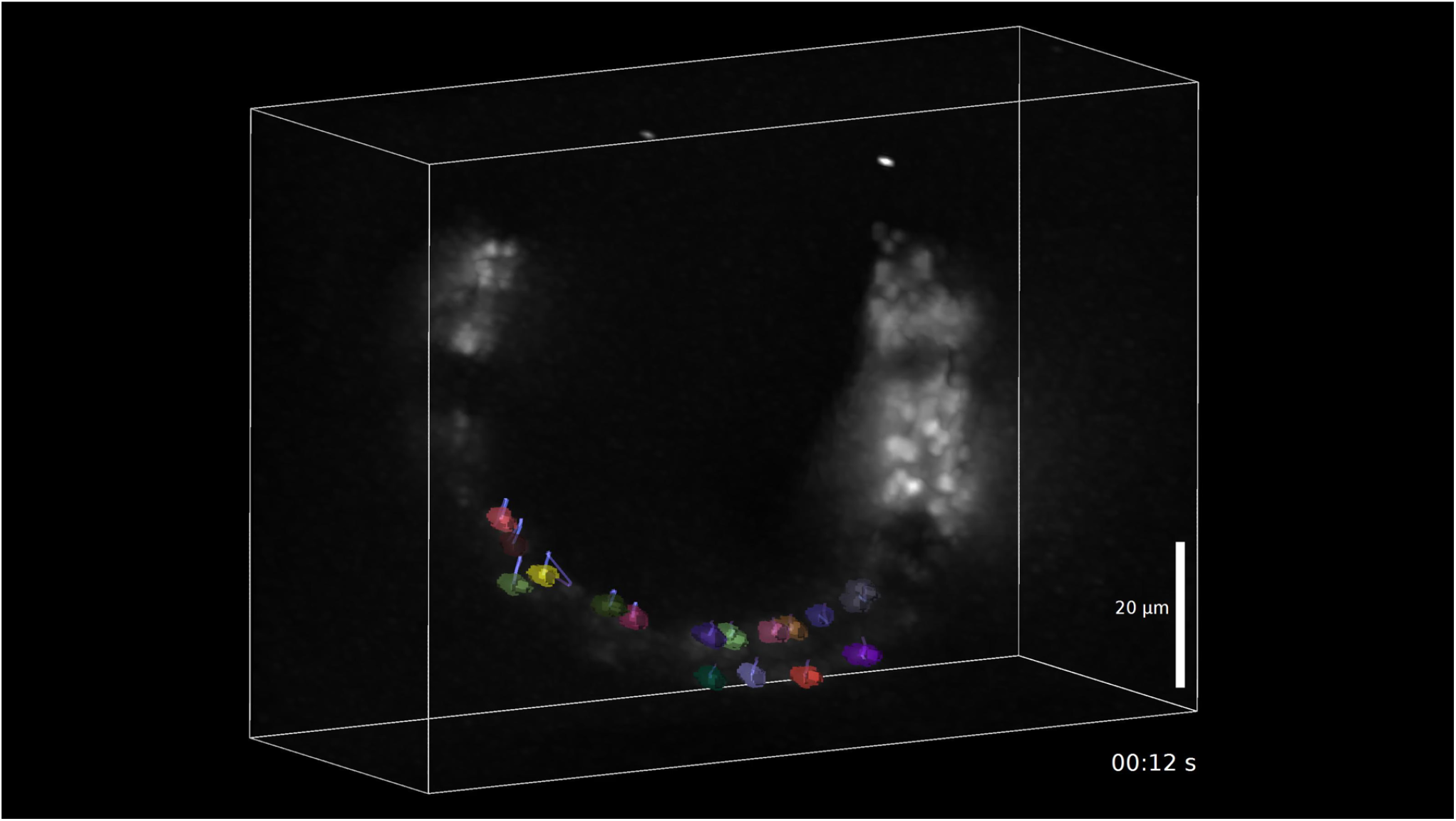
M25 volume showing *C. elegans* (OH1625 strain) with a subset of neurons tracked over time.

**video S7.**
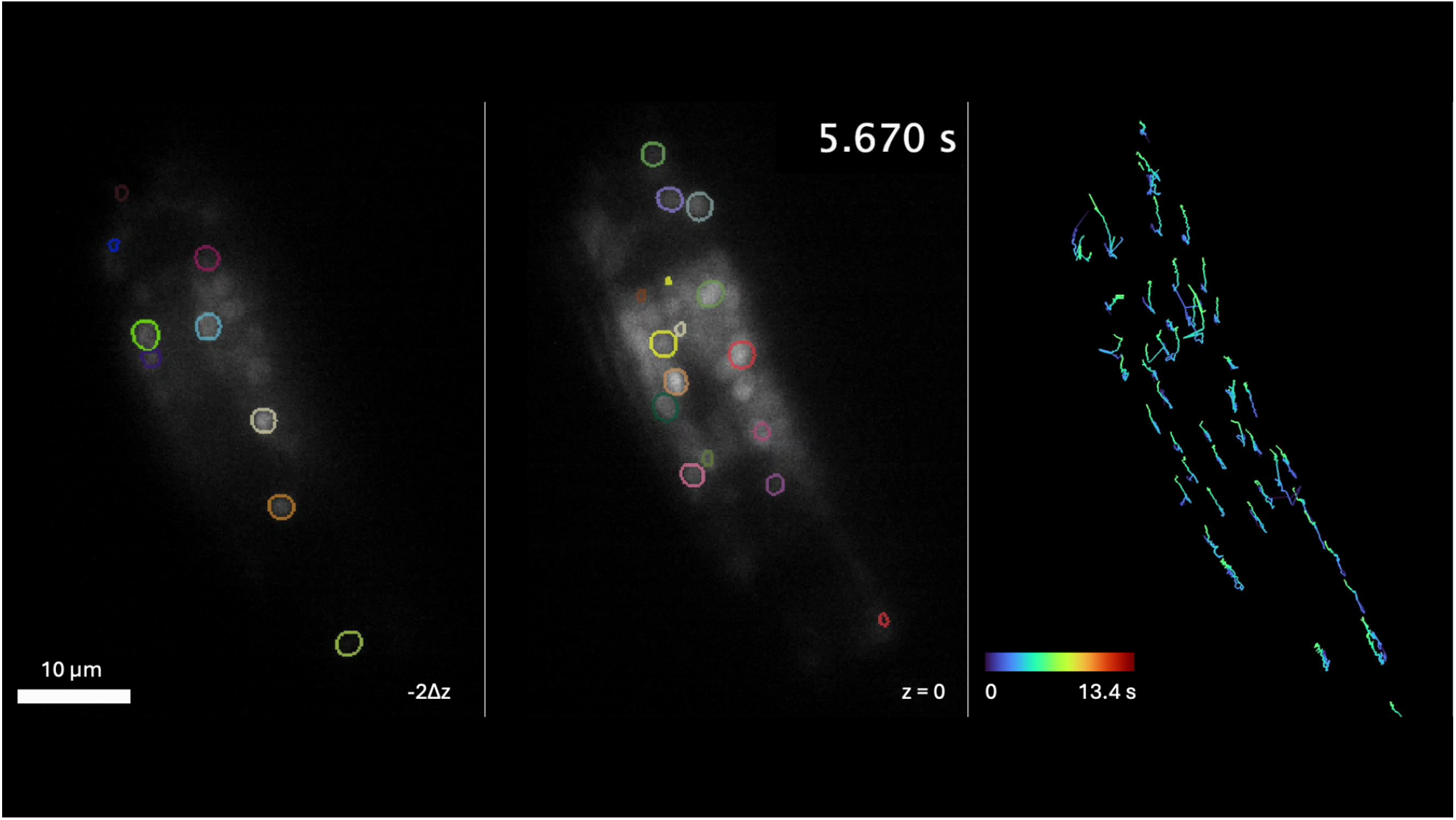
*C. elegans* (OH1625 strain) with a subset of neurons tracked over time at higher magnification.

**Figure S8.**
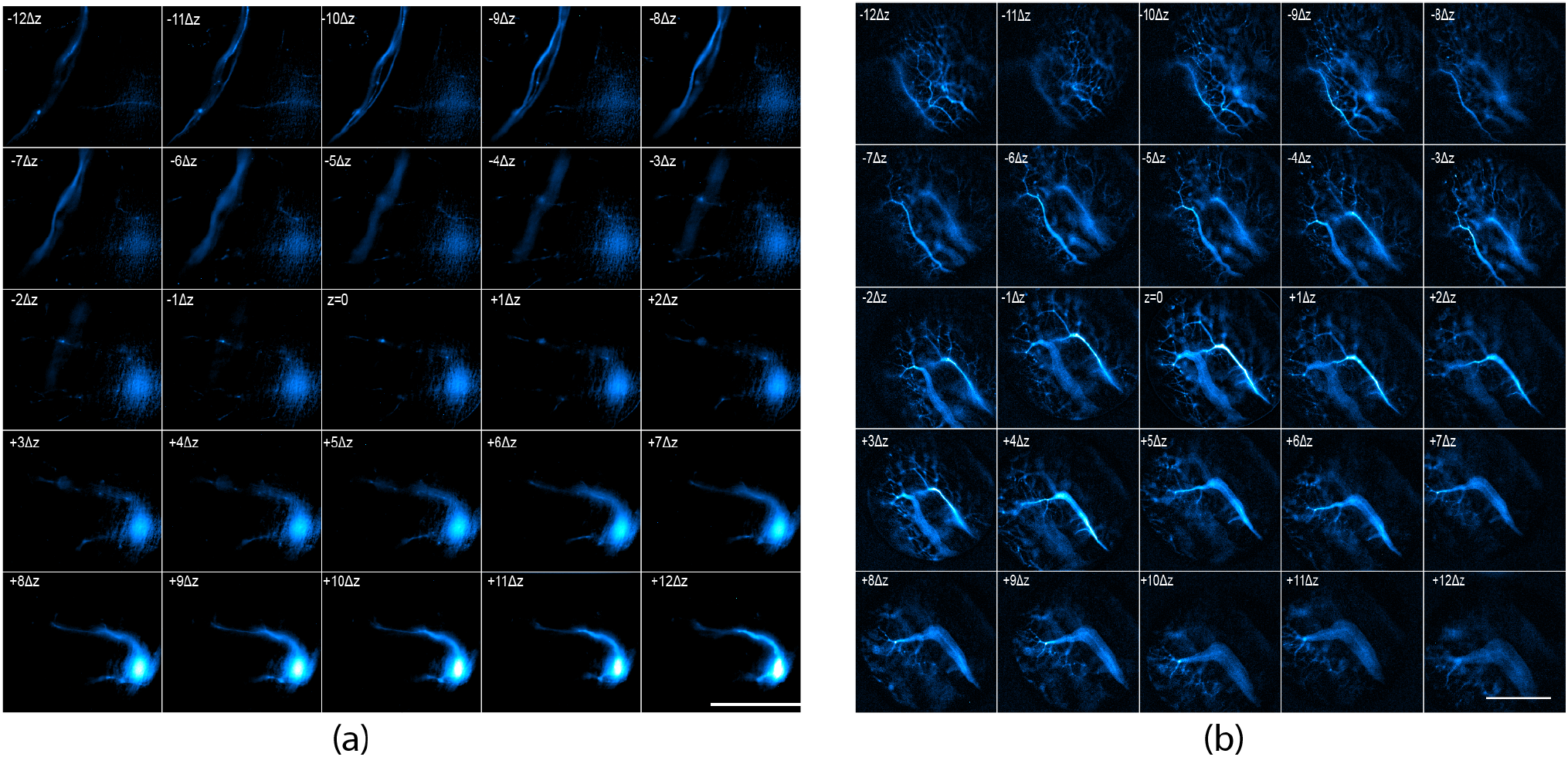
M25 5×5 montage for *P. marinus* spinal cord motor neurons and zebrafish nervous system. (a) Lamprey motor neurons four days post-injection using 10kD Alexa 488. (b) Zebrafish nervous system with endogenous GFP. Used Cyan-hot look-up-table. Scale Bar = 100 µm

**Figure S9.**
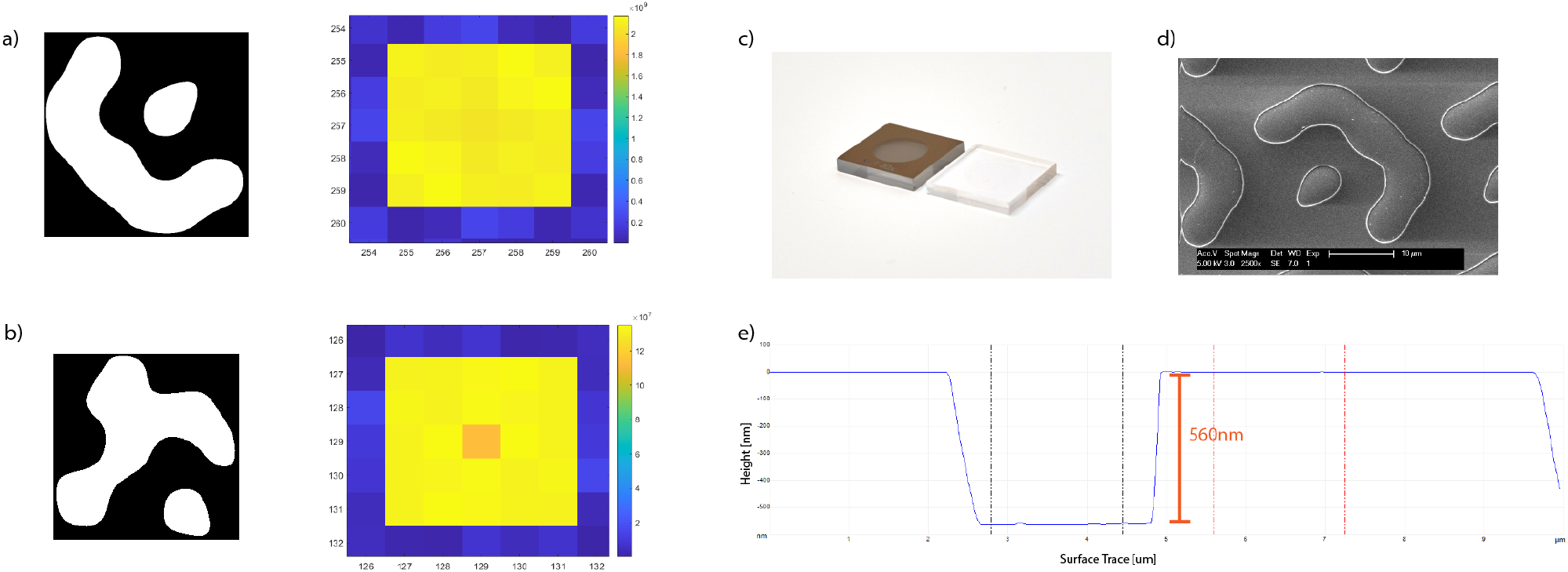
M25 MFG 5×5 Nanofabrication (a) M25 pattern with transmission efficiency of 77% and 0th order suppressed to 96%. Black and white regions represent flat surfaces where one surface is etched down to create a phase shift of *π*. The suppression of the zeroth order *m*_0_(center camera, undiffracted beam) ensures an even signal in the 25 images after the slight light loss in the CCGs, which can be between 5-15% depending on manufacturing quality. (b) M25 pattern with a transmission efficiency of 77% and 0th order suppressed to 86%. (c) Diced MFG with patterned Chrome layer (left) and etched binary phase MFG with the chrome stripped (d) M25 grating function pattern under SEM to confirm etch depth and pattern existence (e) AFM results for M25 MFG dry etch. Precise, sharp walls, and smooth surface etching can be done on quartz photomasks to make custom diffractive optics with the desired phase-shift.

**Figure S10.**
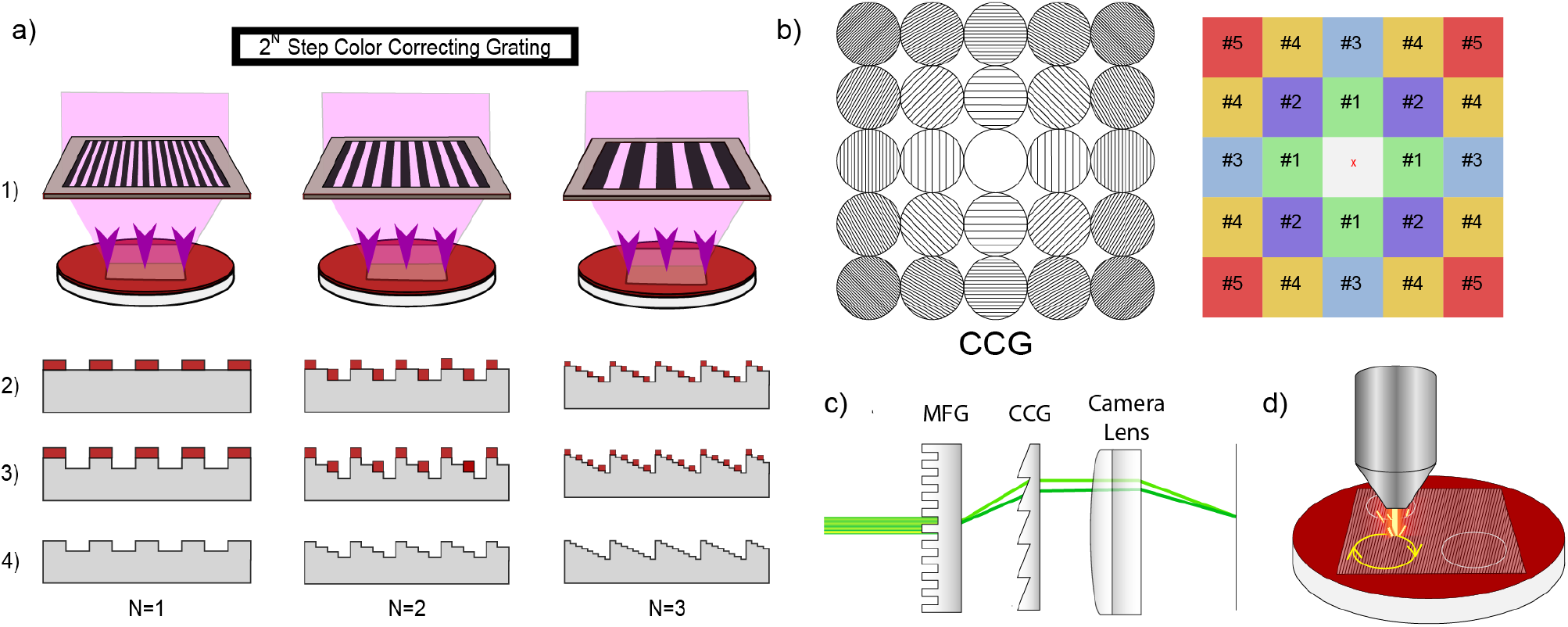
CCG nanofabrication process to create a complete set for the M25. (a) The 3-step photolithography process was used to create an 8-step approximation of the blazed grating, achieving a theoretical transmission efficiency of approximately 90%. (b) Five different patterns are required to correct the chromatic dispersion for each diffractive order, as detailed in Table 2. The 0th order does not require a grating since there is no dispersion along the optical axis. (c) The arrangement of the MFG, CCG, and camera lens demonstrates the new M25 chromatic dispersion correction module. (d) Laser-cutting patterned CCG wafers to produce 29 mm circular gratings for placement in standard camera lens filter holders.

**Figure S11.**
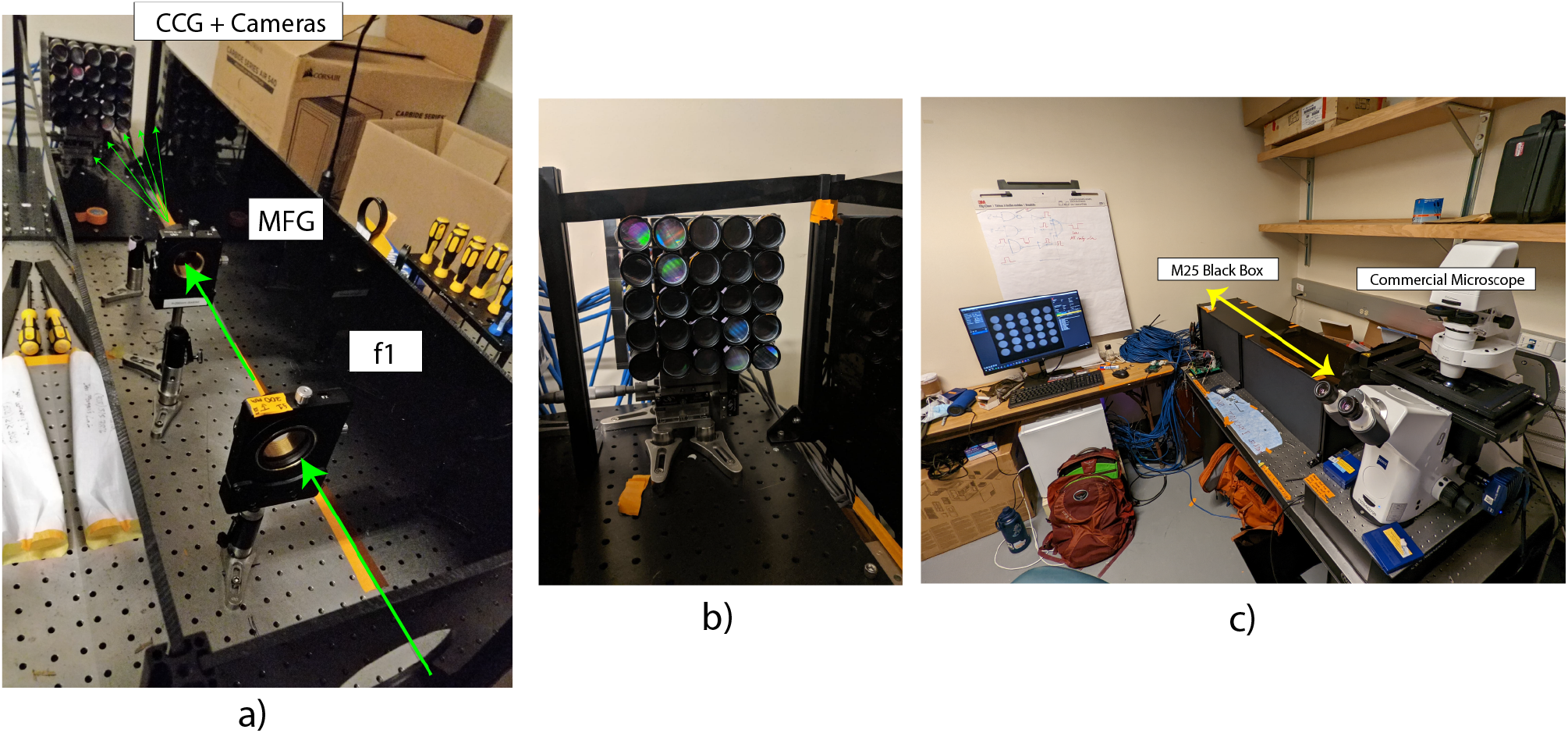
The M25 system can be placed on the side port of any commercial wide-field microscope. (a) Optical path of the beam of fluorescence light coming out of the microscope side-port. Lens f1 forms a Fourier plane where the MFG is placed. The MFG multiplexes and refocuses the beam of light. Each beam is captured on one of the 25 cameras after passing through a CCG and camera lens. (b) 5×5 camera array with a CCG placed in each of the filter holder slots of the lens in the camera array, except for the center camera which captures the zeroth diffraction order and thus does not suffer from dispersion correction. (c) MFM optics are contained in a modular black box at one of the microscope side ports.

